# A transformer-based language model reveals developmental constraint and network complexity during zebrafish embryogenesis

**DOI:** 10.1101/2025.07.09.663853

**Authors:** Juan F Poyatos

## Abstract

Understanding how regulatory complexity and constraint shape organismal development remains a central challenge in biology. The developmental hourglass framework posits that mid-embryogenesis –the phylotypic stage– is a period of heightened conservation and coordinated regulatory organization. We test this hypothesis using Zebraformer, a transformer-based language model trained on single-cell transcriptomic data from zebrafish embryos. Zebraformer learns context-sensitive representations that capture temporal progression, anatomical identity, and regulatory relationships, yielding gene and cell embeddings that recapitulate the developmental axis and increasing transcriptional divergence over time. In contrast, attention-derived gene networks reveal a transient reorganization of regulatory architecture during the phylotypic stage, marked by tightly coordinated gene modules, reduced cross-module connectivity, and diminished local redundancy. Sensitivity to perturbation emerges specifically when regulatory interaction structure is taken into account, rather than from perturbation magnitude alone, highlighting that constraint during this stage is embedded in network topology rather than representational fragility. These findings are supported by graph-theoretic metrics and gene ontology enrichment analyses. Together, our results refine the hourglass framework by localizing developmental constraint to the architecture of gene regulatory networks and demonstrate that language models can extract interpretable biological structure from high-dimensional single-cell data

**Significance statement:** Understanding how cells coordinate to build complex organisms remains a central challenge in biology. Development is genetically encoded yet context dependent, shaped by interactions among genes, cells, and time. Here, we use a transformer-based language model, Zebraformer, trained on single-cell gene expression data from zebrafish embryos to investigate how regulatory organization evolves. The model captures key features of organismal formation, including transcriptional divergence, anatomical specificity, and reorganization of gene regulatory structure. Attention-derived networks reveal a transient phase of tightly coordinated gene modules with reduced cross-module connectivity during the phylotypic stage, followed by specialization. These findings refine the hourglass hypothesis by localizing constraint to network-level organization and demonstrate that contextual language models uncover core principles of biological organization directly from data.

## Introduction

Among the many processes that define living systems, organismal development is perhaps where the contextual nature of biology is most pronounced. By “context,” we refer to the dynamic, situation-specific molecular and cellular states that emerge from both intrinsic genetic programs and extrinsic cues particularly during embryogenesis, when cells differentiate, migrate, and organize into tissues and organs [1].

Technological advances now allow this contextuality to be studied at scale. Single-cell RNA sequencing (RNA-seq) [2], combined with spatial and temporal resolution from highcontent imaging, has opened the door to systematically mapping how gene activity varies across time, space, and cell identity [3, 4]. These tools extend the legacy of classic lineage tracing in organisms such as *Caenorhabditis elegans* [5], offering a view of development as a dynamic, high-dimensional system.

A recent example is the zebrafish (*Danio rerio*) developmental atlas by Lange *et al*. [6], which profiled gene expression at ten developmental timepoints (10 hours post-fertilization [hpf] to 10 days post-fertilization [dpf]) across multiple individuals. This dataset captures a rich temporal trajectory of cellular specification and morphogenesis [7], offering an opportunity to study not just cell fate trajectories, but the broader question of how regulatory structure emerges during development.

Rather than reconstruct individual lineage paths, we ask whether a contextual model trained on this data can reveal emergent principles of developmental constraint and coordination. Specifically, we test whether single-cell expression data contain latent structure that reflect how genes functionally adapt to their cellular environment, and how this structure changes over time. We also explore whether these features can uncover interactions among genes that reflect a dynamically evolving gene regulatory network (GRN) architecture [8].

We place this in the context of the developmental hourglass framework [1, 9, 10], which proposes that early and late embryogenesis are more variable across species, while a midembryonic “phylotypic” stage is highly conserved, marked by a shared body plan and increased developmental constraint. We assess two key predictions of this framework: (1) that the underlying regulatory architecture becomes more complex and centralized during the phylotypic stage, and (2) that this stage is more sensitive to perturbation, reflecting its functional importance.

To test these hypotheses, we introduce Zebraformer, a domain-specific large language model (LLM) adapted from the transformer architecture [11,12]. LLMs, widely used in natural language processing, learn embeddings that capture meaning from surrounding information, an approach increasingly applied to biological systems [13–18]. Zebraformer treats each cell’s transcriptome as a cellular context and learns embeddings for genes based on their activity within that setting. These embeddings encode developmental time, anatomical identity, and functional similarity. Moreover, the model’s attention matrices, originally designed to track relationships among words, offer interpretable proxies for gene-gene regulatory interactions.

We show that this framework captures known biological structure and, critically, offers quantitative evidence for the hourglass hypothesis from a purely data-driven, mechanistically agnostic model. In doing so, we demonstrate that LLMs can serve as hypothesis-generating tools to uncover systems-level organization in development. For background on zebrafish development, the hourglass framework, and transformer-based models in biological contexts, see Supplementary Information.

## Results

### Gene embeddings reflect developmental progression in zebrafish

We began by grouping cells sampled from the same developmental stage and the same sibling fish [6]. For each gene active within these groups, Zebraformer generates a contextual embedding, a *n*-dimensional vector encoding the gene’s behavior in its local transcriptional environment. If a gene’s embeddings are highly similar across cells, this suggests consistent activity or function. To quantify this, we computed all pairwise cosine similarities for each gene’s embeddings across cells, then averaged these scores within fish and across siblings (Fig. 1A, Methods).

**Figure 1:**
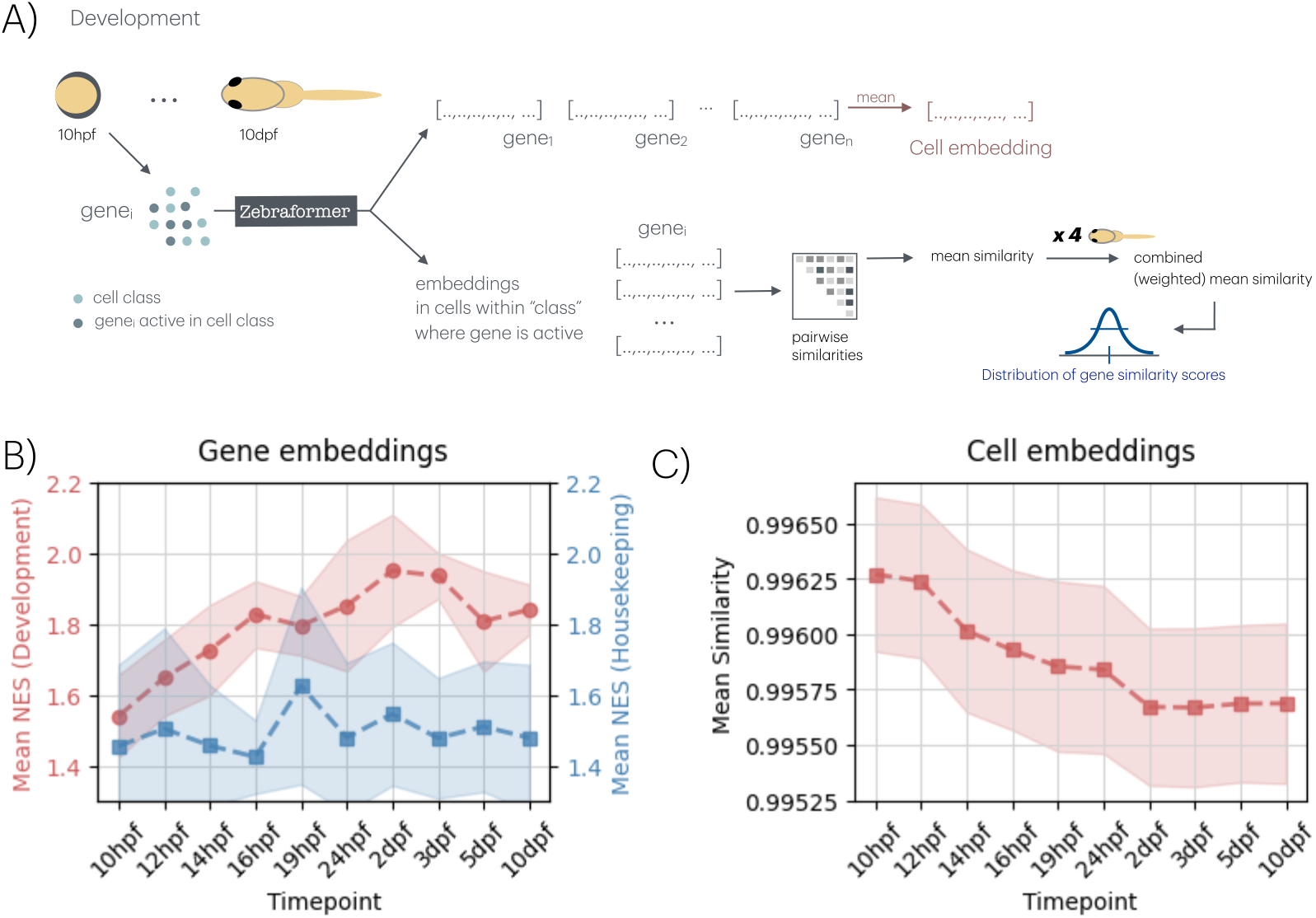
Gene and cell embeddings capture the zebrafish developmental axis. (A) For each developmental stage, Zebraformer generates gene embeddings for cells in a selected class (highlighted). Cell embeddings are computed as the mean of their active gene embeddings. For a gene active in multiple selected cells, pairwise embedding similarities across sibling cells are calculated, then averaged within fish and across siblings, yielding a distribution of similarity scores across genes. (B) Gene Set Enrichment Analysis of genes ranked by embedding similarity across timepoints. Mean normalized enrichment scores (NES) are shown for the top 10 developmental (red) and housekeeping (blue) GO terms. Developmental terms show increasing enrichment over time, while housekeeping terms remain stable. Dots indicate mean NES; shaded areas denote standard deviation (sd). (C) Mean embedding similarity between sibling cells, computed as the cosine similarity of cell embeddings, shaded area indicates sd. Although absolute cosine similarities are compressed near 1, a consistent decrease over time reflects systematic transcriptional divergence and the emergence of cell-type-specific programs.

Because gene embeddings are learned in a high-dimensional space and aggregated across many cellular contexts, cosine similarities are strongly compressed near 1. Accordingly, we focus on relative differences and systematic trends across developmental time rather than on absolute similarity values. To assess which biological processes are associated with consistent gene behavior, we ranked genes by mean similarity score and performed Gene Set Enrichment Analysis. We focused on two functional categories: developmental processes (e.g., differentiation, morphogenesis) and housekeeping functions (e.g., metabolism, transcription). For each category, we selected the top 10 enriched Gene Ontology (GO) terms and plotted normalized enrichment scores (NES) across stages (Fig. 1B, Fig. S1, Methods).

Developmental terms showed steadily increasing enrichment, peaking around 2 dpf, when major anatomical structures are established (Supplement, Table S1). In contrast, housekeeping terms remained stable and low. These results indicate that genes with highly consistent embeddings increasingly correspond to core developmental programs as embryogenesis proceeds, supporting the interpretation that gene embeddings encode biologically meaningful differentiation signals.

### Cell embeddings capture transcriptional divergence during development

To complement gene-level analyses, we examined cell embeddings computed as the mean of active gene embeddings per cell (Fig. 1A). We calculated pairwise similarities between cells at each developmental stage and averaged across sibling fish (Methods). We observed a consistent decrease in average cell-cell embedding similarity over developmental time (Fig. 1C).

Although the absolute change in mean cosine similarity is numerically small, this trend is robust across individuals and reflects a systematic restructuring of embedding geometry rather than stochastic variation. In this compressed similarity regime, such shifts correspond to increasing transcriptional divergence as cells acquire distinct identities. This interpretation aligns with principles of developmental biology and reinforces the idea that Zebraformer embeddings capture progressive differentiation rather than static cell identity.

### Embedding similarity varies across anatomical domains

We next asked whether embedding similarity patterns are conserved within specific anatomical regions. Using curated domain annotations (e.g., lateral mesoderm, neural crest, hematopoietic system; domains in Zebrahub refer to *embryonic tissues* or *cell lineages* in zebrafish and vertebrates generally [19], Supplement, Table S1), we computed *per*-gene embedding similarities across cells within each domain, stage, and fish. Trends were analyzed across timepoints.

Figure 2 highlights three representative cases. The lateral mesoderm (LM) and neural crest (NC) show decreasing embedding similarity over time, whereas the hematopoietic system (HS) shows an increasing trend. We emphasize that these examples are chosen to illustrate contrasting developmental dynamics rather than to suggest that all domains exhibit strong temporal trends. Indeed, some of them display relatively flat trajectories (Fig. S2), which is expected given the diversity of developmental programs captured by the annotations.

**Figure 2:**
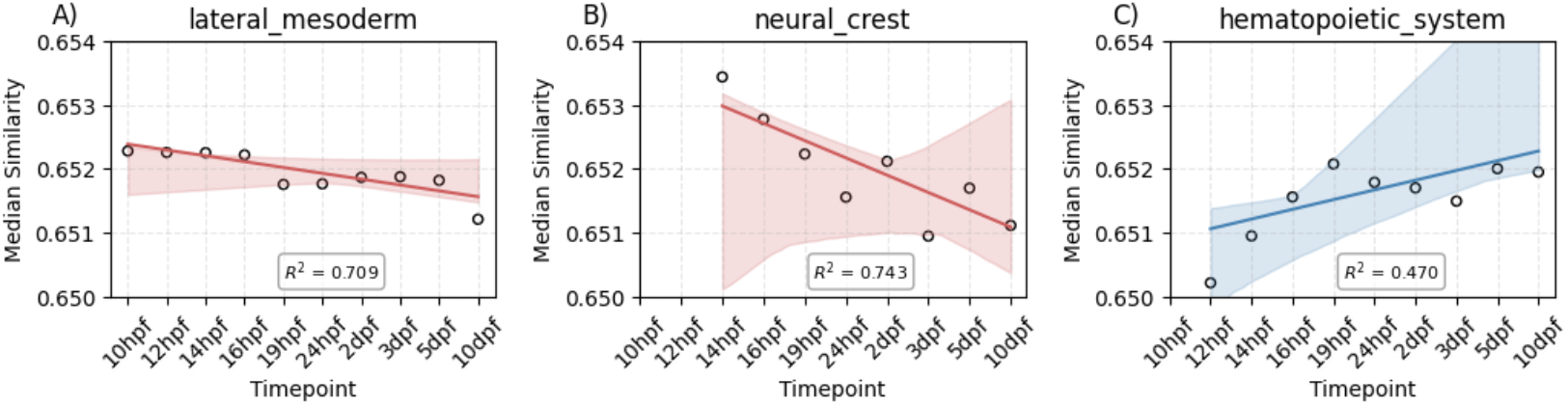
Gene embeddings reflect anatomical context and temporal progression. For each anatomical class, we plot regression trends of median gene embedding similarity over time. Similarity is computed *per* gene across cells in which it is active, then summarized at the domain level. Shown are three representative domains illustrating distinct developmental dynamics: (A) Lateral mesoderm, where similarity decreases with time (*R*^2^ = 0.709); (B) Neural crest, showing a stronger decrease with higher uncertainty (*R*^2^ = 0.743); and (C) Hematopoietic system, where similarity increases over time (*R*^2^ = 0.470). Shaded areas indicate bootstrapped uncertainty from 10,000 resamples.

These patterns align with known biology. In zebrafish, the LM, including the lateral plate mesoderm, undergoes extensive fate diversification and spatial reorganization from early somite stages through larval development [20, 21]. Similarly, the NC originates as a highly multipotent population and gives rise to diverse derivatives, including neurons, cartilage, and pigment cells, following strongly context-dependent spatiotemporal trajectories [22]. Increasing functional specialization in these domains is therefore expected to reduce average embedding similarity over time. In contrast, the HS follows a more hierarchical, tree-like developmental program in which hematopoietic stem cells maintain multipotency through self-renewal while gradually restricting lineage potential [23], consistent with increasing embedding similarity.

Importantly, the coexistence of decreasing, increasing, and flat trends across domains indicates that Zebraformer does not impose uniform temporal structure, but instead captures biologically meaningful heterogeneity in developmental dynamics across tissues and lineages.

### Contextual diversity increases in late-emerging cell types

To examine gene contextuality at finer resolution, we analyzed 154 annotated cell types across timepoints. For each cell type and stage, we computed median gene embedding similarity across cells, averaged over sibling fish (Methods). This metric reflects how consistently genes behave in a given cell type at a given time.

Strikingly, cell types with lower average similarity tended to appear later in development, while those with higher similarity emerged earlier (Fig. 3, Fig. S3). Early cell types are likely defined by more generic transcriptional programs, involving reused “foundational” genes. Later-emerging cell types express more specialized or spatially restricted gene sets, resulting in more diverse embedding patterns. This finding reinforces the idea that embedding similarity serves as a proxy for transcriptional specialization.

**Figure 3:**
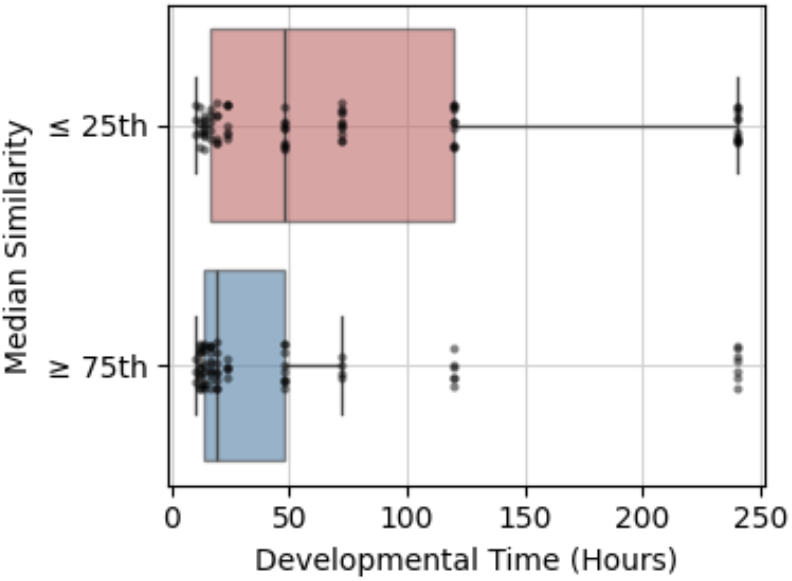
Cell types with lower average gene embedding similarity tend to emerge later in development. For each cell type at each timepoint, we compute the median embedding similarity of active genes across individuals. Cell types are grouped into quartiles based on similarity. The box plot shows developmental timing (hours post-fertilization) for high-similarity (≥ 75th percentile) and low-similarity (≤ 25th percentile) groups. Cell types with lower embedding similarity emerge significantly later in development (Mann–Whitney U test, *p* = 0.00676).

### Attention-based networks reveal hourglass structure of development

Can we go beyond the relational information captured by embeddings? Zebraformer’s attention matrices offer another layer of information, capturing gene-gene associations learned during training [12]. These attention-derived graphs can be interpreted as datadriven GRNs, allowing us to assess regulatory architecture across developmental time.

We use this tool to assess how well Zebraformer captures the hourglass framework of development [1, 9, 10] (Supplement). We divided development into three stages: early (≤ 14 hpf), middle/phylotypic (15–19 hpf), and late (≥ 24 hpf) [24]. For each, we built attention-based GRNs. To avoid trivial effects driven by node set expansion and graph density, we constructed these networks on a fixed gene universe, enforced controlled sparsity (top 5 targets per gene), and normalized edge weights by source gene frequency. Under these conditions, network size and density increase monotonically from early to late development (Fig. S4), reflecting increasing transcriptional diversity, but these global trends alone do not account for stage-specific organization (Methods).

Community structure analysis revealed that modularity remains comparable between early and phylotypic stages but decreases at later stages, indicating a loss of modular organization during post-phylotypic development. In contrast, the phylotypic network shows a pronounced reduction in cross-module connectivity: the fraction of total edge weight linking genes assigned to different communities reaches a minimum at the phylotypic stage (Fig. 4A). This reduction in inter-module coupling is accompanied by a decrease in the clustering coefficient, indicating decreased local redundancy and fewer triadic interactions among genes. Power-law analysis of the degree distribution reveals a modest peak in the scaling exponent *α* at the phylotypic stage (Fig. 4B), indicating a transient reduction in extreme hub dominance relative to early and late stages (Supplement, Table S2).

**Figure 4:**
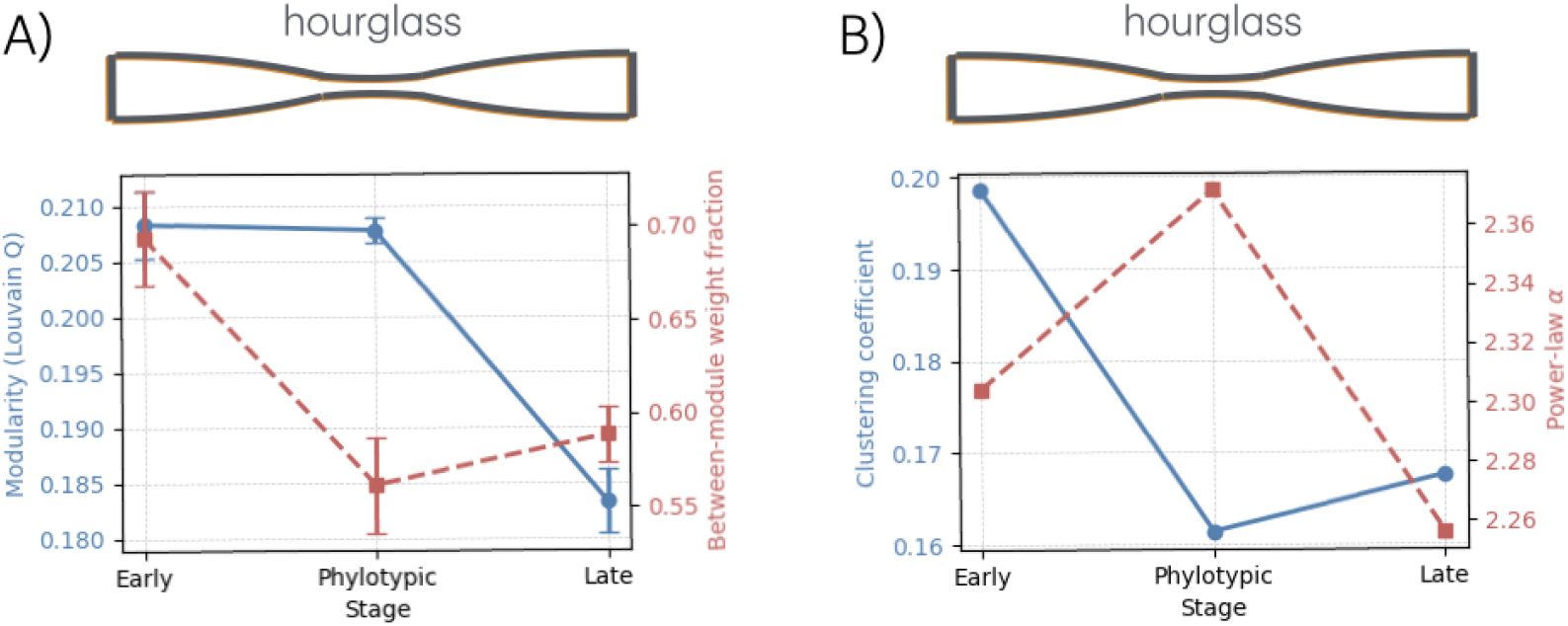
Attention-based gene regulatory networks exhibit stage-specific organization consistent with the hourglass framework. (A) Louvain modularity (blue, left y-axis; mean ± sd across 20 random initializations) is comparable between early and phylotypic stages but decreases at later stages. In contrast, cross-module coupling, the fraction of total edge weight connecting genes assigned to different communities, reaches a minimum at the phylotypic stage, indicating reduced inter-module connectivity (red, right y-axis; mean ± sd). (B) The clustering coefficient (blue, left y-axis) shows a minimum at the phylotypic stage, consistent with reduced local redundancy, while the power-law exponent *α* (red, right y-axis) exhibits a peak at this stage, indicating a transient reduction in extreme hub dominance. The hourglass schematic summarizes the classical expectation of a mid-embryonic constriction in developmental variability.

These features identify the phylotypic stage as a distinct regime characterized by tightly segregated, internally coordinated gene modules with limited cross-talk. Early developmental networks exhibit higher inter-module connectivity and greater local redundancy, consistent with a more flexible and exploratory organization, whereas late developmental networks show increased connectivity but reduced modular structure. Thus, rather than reflecting maximal global interactivity, the developmental waist corresponds to a phase of heightened modular specialization and reduced redundancy, consistent with an hourglass-like pattern of constraint arising from tight coordination within essential developmental programs.

### Topology-aware perturbations reveal constrained interaction structure without increased embedding fragility

The hourglass framework predicts heightened vulnerability to perturbation during the phylotypic stage, attributed to dense regulatory coupling and global coordination of developmental programs [25]. To test this prediction, we assessed how gene embeddings learned by Zebraformer respond to increasingly structured *in silico* perturbations across developmental time.

Initial experiments using random perturbations, ranging from single-gene masking to stronger, multi-gene ablations, produced relatively uniform embedding-level responses across pre-phylotypic, phylotypic, and post-phylotypic stages. These included strategies targeting phylotypic-enriched transcription factors and eligibility-matched multi-hit designs. Across all such conditions, we observed no clear peak in phylotypic embedding displacement (Supplement, Tables S3-S6, Fig. S5).

To specifically examine whether developmental sensitivity emerges from interaction structure, we implemented topology-aware perturbations derived from attention-based GRNs. At each stage, hub genes were identified by node degree, and fixed-size neighborhoods around these hubs were defined. Entire hub-centered neighborhoods were then masked concurrently, simulating disruption of local regulatory modules. Perturbation impact was quantified as the resulting shift in cell embedding space, compared against randomized controls that masked genes of similar degree but without structured connectivity.

Hub-centered perturbations consistently induced stronger embedding displacements than their degree-matched controls confirming the presence of non-random, topologically meaningful dependencies. However, the magnitude of this effect was similar in the pre-phylotypic and phylotypic stages, and notably reduced in the post-phylotypic stage (Fig. 5, top).

**Figure 5:**
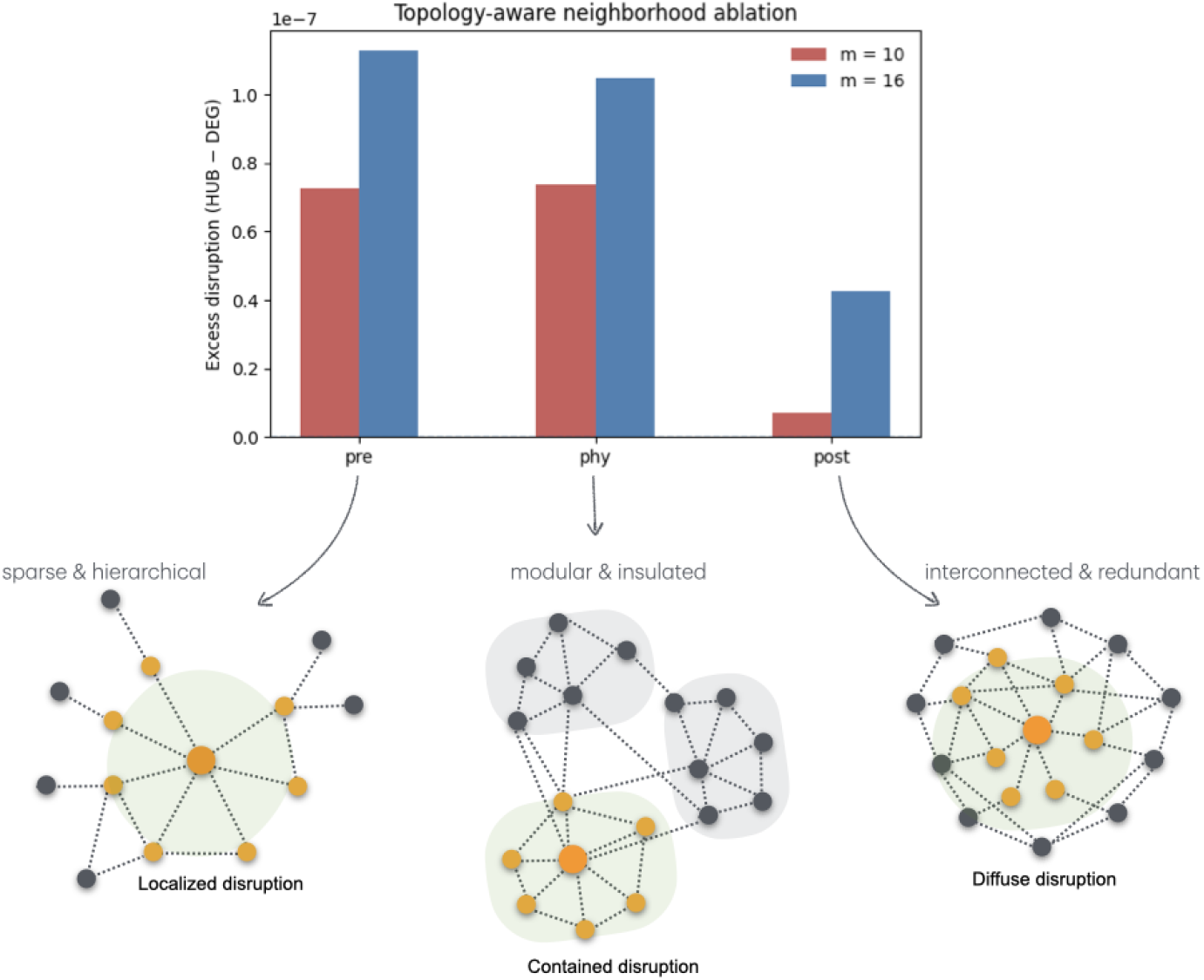
Topology-aware neighborhood ablation reveals distinct disruption regimes across stages. Fixed-size neighborhoods (*m* = 10, 16) centered on high-degree genes (‘hubs’) were ablated in attention-derived GRNs across developmental stages. Disruption, quantified as the mean aligned shift in non-ablated gene embeddings, was compared to degree-matched random controls. Bars show excess disruption attributable to network topology. Effects were comparable in pre- and phylotypic stages and reduced post-phylotypically. Schematic panels below illustrate disruption modes: nodes are genes, dashed lines denote interactions, orange hubs mark ablated genes, and green shading indicates affected regions. Pre-phylotypic networks are sparse and hierarchical (localized disruption); phylotypic networks are modular and insulated (contained disruption); post-phylotypic networks are dense and redundant (diffuse disruption). These patterns reflect changing topological regimes underlying developmental robustness (see also Fig. 4).

This pattern can be understood by considering stage-specific network features. Although modularity remains relatively stable from early to phylotypic stages, cross-module connectivity drops significantly at the phylotypic stage (Fig. 4A). This suggests that during mid-development, gene-gene dependencies become more localized within modules and less distributed across the network. Under these conditions, fixed-size neighborhood ablations at hubs may disrupt similar numbers of within-module connections in both early and phylotypic stages, leading to comparable sensitivity. In contrast, the late-stage network exhibits lower modularity and higher redundancy, buffering the effect of local ablations (Fig. 5, bottom).

Together, these findings reveal that Zebraformer embeddings are robust to unstructured perturbations but sensitive to organized disruptions that reflect regulatory architecture. However, this sensitivity does not peak at the phylotypic stage in embedding space. Rather, the developmental hourglass emerges in the topology of GRNs, with mid-stage development exhibiting maximal modular separation and reduced cross-talk. Constraint, in this view, is encoded not in representational instability, but in the global structure of regulatory dependencies.

## Discussion

Development is an inherently contextual, multiscale process. Our goal was to ask whether a language model –trained not to predict but to represent– could capture the regulatory logic that underpins this process, particularly the constraints that emerge during key developmental transitions. Using Zebraformer, we show that gene embeddings learned from single-cell expression data reflect both developmental time and anatomical context. Early in development, genes exhibit high embedding similarity across cells, consistent with broad, shared transcriptional programs. As development proceeds, these representations diverge, especially in later-emerging, specialized cell types. This supports the idea that Zebraformer learns not just expression levels, but context-dependent regulatory roles.

Beyond representation, the model’s attention matrices, used here as proxies for GRNs, reveal a striking temporal pattern consistent with the hourglass concept of embryogenesis [1,9, 10]. While overall network size and density increase monotonically with developmental time, stage-specific differences emerge in higher-order organization. In particular, the phylotypic stage is characterized by reduced inter-module coupling, decreased local redundancy, and a transient shift in degree organization, resulting in tightly segregated yet internally coordinated gene modules. Early developmental networks exhibit greater cross-module connectivity and redundancy, whereas later networks show increased connectivity accompanied by reduced modular structure. Together, these features identify the developmental waist as a phase of heightened modular specialization rather than maximal global interactivity.

Perturbation analyses clarify how this organizational constraint is expressed. Despite pronounced stage-specific reorganization at the network level, embedding representations remain remarkably robust across development. Neither single-gene perturbations nor structured, topology-aware neighborhood ablations reveal a phylotypic peak in embedding sensitivity. Even when perturbations respect interaction structure by targeting high-degree hub neighborhoods, embedding disruptions are comparable between pre-phylotypic and phylotypic stages and attenuated later in development. These findings indicate a separation between representation and interaction: while attention-based networks encode stage-specific regulatory constraint, the geometry of learned embeddings remains smooth and redundant. Developmental constraint therefore manifests primarily through the organization of gene-gene dependencies, rather than as increased fragility of embedding representations.

Importantly, our results demonstrate that sensitivity to perturbation emerges only when regulatory interaction structure is taken into account, not simply from perturbation magnitude alone. This reinforces the idea that the phylotypic stage is defined by global coordination and interdependence of regulatory modules, even as the embeddings themselves remain stable. The decoupling of representational stability and network fragility offers a refined understanding of constraint: it is not the expression space that becomes brittle, but the structure of influence and dependency.

Our attention-based network analysis reveals stage-specific transitions in regulatory influence. When this is quantified using normalized, sparsified, and directed attention networks, genes that predominantly receive regulatory control early in development are enriched for transcriptional and chromatin-associated functions, indicating convergence onto a conserved regulatory substrate. At the phylotypic stage, regulation is most strongly focused on chromatin- and nucleosome-related processes, consistent with centralized coordination of developmental programs. In contrast, later stages are characterized by genes that predominantly exert influence and are enriched for signaling, receptor, and structural functions, reflecting the emergence of specialized and tissue-specific regulatory control (Supplement, Table S7). This progressive redistribution of regulatory influence is consistent with models of developmental canalization, in which early and mid-developmental constraints shape downstream trajectories while permitting diversification at later stages [26, 27].

Unlike foundational models trained across multiple tissues, species, or experimental contexts (e.g., scBERT [13], Geneformer [14,15], scGPT [16], and scFoundation [17]; Supplement, Table S8), Zebraformer is deliberately scoped to zebrafish embryogenesis. While this limits generalizability, it enables finer resolution and more interpretable embeddings, revealing patterns that might be diluted in broader corpora. Our aim is not universality, but biological clarity, a model that reflects the temporal and contextual logic of development.

More broadly, this work highlights the potential of LLMs as hypothesis-generating frameworks in biology. Rather than predicting annotations or clustering cells, we use the model to infer structure, both in the gene embedding space and in emergent networks. This mirrors a shift in the field, where transformer-based models are increasingly applied to uncover relational structure in biological systems [28]. Intriguingly, the attention mechanism itself may reflect a computation analogous to that of biological networks, context-aware, nonlocal, and dynamically weighted [29].

In sum, our findings demonstrate that a transformer model trained purely on expression context can recapitulate key features of developmental constraint, supporting long-standing theoretical models from a new, data-driven angle. As models like Zebraformer become more integrated with experimental approaches, we anticipate that contextual language models will play a growing role in developmental and systems biology not as black-box predictors, but as interpretable lenses onto the organizational logic of living systems.

## Methods

### Large language model

We apply masked language modeling (MLM) following the Geneformer approach [14], see also [13]. To this aim, we use the BertForMaskedLM model from Hugging Face’s Transformers library [30], a widely adopted toolkit that provides access to pretrained transformer models. BertForMaskedLM is a variant of BERT [31] fine-tuned for MLM, i.e., predicting missing tokens in input text (Supplement, Figs. S6–S8).

### Developmental dataset

We downloaded the single cell RNA-seq data of the zebrafish developmental trajectory from Zebrahub (Supplement). We retain cells with at least 100 expressed genes and genes detected in at least 3 cells, excluding those with extensive mitochondrial or noncoding expression. This results in a final dataset of 111,249 cells (acting as “sentences”) and 28,677 genes (that will be part of the “tokens”, the small units that serve as the input for the LLM).

### Rank value encoding and tokenization

Each gene’s activity, as captured by the singlecell transcriptome, is represented by a rank value. Genes are ranked by their expression level in each cell normalized by the non-zero median expression of each gene across the full developmental dataset. The protocol implies that specific genes, e.g., transcription factors, and housekeeping genes receive higher or lower prioritization in the rank-based encoding, respectively [14]. This ordering is analogous to the word order in a sentence and represents the input provided to the LLM after tokenization. Tokenization is done using a dictionary of length 28,679 (28,677 genes and two special tokens: ⟨pad⟩ as 0 and ⟨mask⟩ as 1). The number of genes per cell is limited to a maximum sequence length of 512 tokens to control computational cost (sequence length is 2,048 in Geneformer [14] and 512 in standard BERT [31]).

### Zebraformer training

We used Optuna, an open-source Python library for hyperparameter optimization [32], to jointly tune architectural and training hyperparameters. The selected model has an embedding dimension of 384, 2 transformer layers, 3 attention heads, and a maximum input length of 512 tokens. The feed-forward network intermediate size was set to 1152 (3× the embedding dimension). Input and output embeddings are tied, so the masked-token prediction head reuses the input embedding matrix rather than introducing a second vocabulary-sized parameter matrix. Model capacity is therefore determined by the vocabulary size, embedding dimension, and the number and width of transformer layers, resulting in a model with on the order of 10^7^ trainable parameters. Training batch size and learning rate were chosen to accommodate GPU memory constraints. We used the AdamW optimizer with weight decay, and the final pretrained model achieved nearly identical MLM losses on training and validation sets, indicating no overfitting despite the modest corpus size.

### Gene and cell embeddings

We select the penultimate hidden layer of the model to extract *gene* embeddings. While the final layer is typically more closely tied to the model’s training objective, the penultimate layer tends to be more representation-focused, capturing contextual information in a way that is more transferable across tasks [33]. To compute a representative embedding for each *cell*, we aggregated gene-level embeddings across all genes active in the particular cell, normalized by the number of contributing genes. This yielded a mean embedding vector for each cell that reflects the average of its active gene embeddings (Fig. 1A).

### Weighted gene similarities

For each gene *i*, we computed a similarity score based on the cellular contexts in which it is active and the broad class of cells considered. Specifically, we defined three broad classes based on the following groupings: (i) by timepoint alone, (ii) by anatomical domain within a specific timepoint, or (iii) by cell type within a specific timepoint. Within each class, we further partitioned cells by sibling fish identity. For each sibling group *k*, we computed the mean pairwise *cosine* similarity *µ*_*i*_ across all pairs of gene embeddings **e**_*a*_ and **e**_*b*_, for all cell pairs *a* and *b* in the group, where similarity is defined as 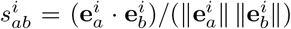 We then calculated the weighted average similarity across all sibling groups as 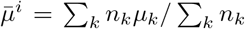, where *n*_*k*_ is the number of contributing cells in each group, and ∑_*k*_ *n*_*k*_ gives the total number of contributing cells (Fig. 1A). To ensure robust estimates, we included only genes that were active in more than four cells (on average) in at least three different fish, and had a pooled standard deviation *s*_pooled_ *<* 0.05, computed by combining within-fish variance and between-fish deviation from the overall mean. Moreover, to quantify the average of all gene similarity scores, we alternatively use the median as a descriptor of the distribution when they are not normal (Figs. 1–2).

### Weighted cell similarities

To quantify transcriptional consistency within individual fish over developmental time, we computed pairwise cosine similarity scores between cell embeddings for each fish and timepoint. For each class, we recorded both the number of cells and the mean pairwise similarity. As in the gene-level analysis, we aggregated *per*-fish statistics by computing a weighted average similarity. These aggregated values enable longitudinal comparisons across developmental stages.

### Normalized enrichment score

We perform gene set enrichment analysis to assess whether gene ontology (GO) terms related to development and housekeeping functions are associated with stronger gene embedding similarity as time progresses. To this end, we rank the gene similarity scores for each timepoint and test whether a given biological process term (GO Biological Process 2021 database) is concentrated near the top (strong similarity) or bottom (weak similarity) of the list using the official Python 3 interface to the g:Profiler toolkit [34]. This is quantified by the normalized enrichment score (NES), with positive values indicating enrichment. We select specific developmental or housekeeping terms (Fig. S1) and compute their mean NES and standard deviation of the top 10 instances. These values are shown in Fig. 1B.

### Similarities and anatomical domais

Each gene can be active in a number of cells associated with an anatomical term (e.g., the Jun dimerization protein 2b, jdp2b, is active in the central nervous system related cells at timepoint 10hpf). We take the mean and sd of the gene embeddings similarities on such set and compute again a combined (weighted) mean and pooled sd as before (considering the number of cells on each fish; and the filtering for robust estimates discussed above). For each anatomical domain, we calculate a robust trend analysis using Theil–Sen regression by considering timepoints (categorical timepoints encoded as integers, e.g., “24hpf” → 0, “48hpf” → 1, etc.,) and the associated median similarity values for those timepoints.

### Attention matrices

We computed attention scores for each of the three developmental stages described by the hourglass framework. The early stage includes data from cells collected at 10hpf to 14hpf; the phylotypic (middle) stage includes 16hpf and 19hpf; and the late stage includes 24hpf to 10dpf. For each stage, attention scores were aggregated across all cells and attention heads, with edge weights proportional to the raw attention magnitudes to preserve interaction strength. Outgoing weights were normalized by the number of occurrences of each source gene to prevent highly prevalent genes from dominating the network. To ensure comparability across stages, we applied controlled sparsification by retaining only the top 5 strongest outgoing edges per gene, yielding networks with matched density and node sets. The resulting graphs therefore capture differences in the consistency and organization of gene-gene coordination rather than effects of sampling depth or gene frequency, and were used for downstream topological analyses (e.g., modularity).

### In silico perturbation

We performed a series of *in silico* perturbation analyses in which genes were systematically masked from cell-specific ranked gene sequences and embedding responses were recomputed using the pretrained model. Perturbation impact was quantified as the mean cosine distance between wild-type and perturbed embeddings across all non-masked positions, and per-cell sensitivity scores were obtained by averaging across multiple perturbations. Perturbation schemes ranged from isolated single-gene masking to coordinated multi-gene perturbations and topology-aware perturbations that explicitly respect attention-derived gene interaction structure. To ensure fair comparisons across developmental stages, perturbations were restricted to shared gene panels, normalized for gene availability where appropriate, and evaluated on matched cell populations. Detailed descriptions of perturbation protocols, control analyses, and additional sensitivity metrics are provided in the Supplement.

## Acknowledgements

This work was supported by grant PID2023-151289NB-I00 funded by MICIU/AEI/10.13039/501100011033 and by “ERDF/EU”. We thank Joaquin Torres for discussions on LLMs, Merlin Lange and Yang-Joon Kim for assistance with the Zebrahub dataset, and the Computational Systems Biology Group at the Spanish National Biotechnology Center for providing access to their computational resources. This work was done with a single GPU unit.

## Code availability

Code available at github.com/juanfpoyatos/Zebraformer. The author used OpenAI’s Chat-GPT to assist with text improvement and code debugging. The author reviewed and edited all content and is fully responsible for the final manuscript.

