## Supplement for "A transformer-based language model reveals developmental constraint and network complexity during zebrafish embryogenesis"

Juan F Poyatos

National Museum of Natural Sciences (MNCN-CSIC), Madrid 28006, Spain.

We introduce below the supplementary figures along with the key concepts, tools, and datasets used throughout this manuscript, aiming to present them in a concise and accessible manner for readers unfamiliar with the topics. More detailed discussions are available in the referenced literature. We also include further analysis associated with the discussion in the main text.

#### The hourglass model of embryogenesis

During embryogenesis, the first cell formed when a sperm fertilizes an egg develops into a complex, differentiated adult organism by passing through a series of distinct stages and forming specific morphological structures along the way. Embryologists such as Karl Ernst von Baer, who proposed that development proceeds from general to specific features [1], and Ernst Haeckel, who introduced the (now discredited) idea that an organism's development (ontogeny) recapitulates its evolutionary history (phylogeny) [2], laid the foundation for early hypotheses about how these structures emerge and the mechanisms that drive their formation.

Building on these early ideas, later researchers proposed more refined models of developmental processes and their conservation across species. Notably, Duboule argued that development follows a developmental hourglass model [3, 4] (see Fig. 4B, main text), a concept also supported by Raff [5]. In this model, early and late stages of development are less evolutionarily conserved, while a middle period is highly conserved among different organisms. It is during this stage that the common anatomical features of the basic body plan are established. Because ancient organismal lineages (called *phyla*) differ in their body plans, this is why the middle phase is known as the *phylotypic* stage.

In early development, germ layers (three simple layers of cells –ectoderm, mesoderm, and endoderm) and the major body axes and tissue-specific domains of the embryo are established. These processes are controlled by relatively simple and modular gene regulatory networks (GRNs), the set of interactions showing how genes control each other, with some genes turning others `on` or `off` to drive specific cellular actions [6,7]. During the middle stage, organogenesis takes place, and GRNs reach peak complexity and evolutionary constraint; even small changes at this point could lead to severe developmental defects [8], forming the narrow waist of the hourglass model. In late development, limbs and organs mature, GRNs become more specialized and compartmentalized, and changes in gene activity tend to have less impact. See [9] for a broader discussion.

#### The `ZebraHub` dataset

`Zebrahub` is a cutting-edge resource that maps how cells differentiate in zebrafish embryos over time. It integrates detailed time-lapse videos of living embryos with information on which genes are active or inactive in each individual cell as they move and gradually take on specific roles in the body. Using advanced tools like light-sheet microscopy, to capture clear images of embryos with minimal damage, and single-cell RNA sequencing (scRNA-seq), which reveals what each cell is doing at the genetic level, `Zebrahub` provides a rich, layered view of how development unfolds in both space and time. The imaging captures the entire embryo at once, a method known as *in toto* imaging, so no part is missed as cells divide, shift position, and specialize.

Note that to build these cell maps, we can use different strategies. One is to combine cells from several embryos of the same age (called pooled embryos) to increase the number of cells analyzed, though this approach assumes embryos are all developing in the same way. Another method uses multiplexed embryos, where each embryo is labeled with a unique marker, making it possible to study individual variation but requiring more complex experiments and analysis. Importantly, `Zebrahub` also studied sibling embryos, i.e., those from the same parents, at different points in development, offering a more consistent and controlled look at how embryos grow. Note also that biological *domains* (cell types, tissues, lineage labels) are curated through manual annotation of clusters using marker genes and external knowledge-based ontologies.

More information available here https://zebrahub.sf.czbiohub.org and in the corresponding manuscript [10]. Additional scRNA-seq studies have provided valuable insights into zebrafish development up to 24 hpf, e.g., [11–15].

#### Zebrafish developmental timepoints and anatomical domains

The following terms, available as annotation in the `ZebraHub` dataset, refer to embryonic tissues or cell lineages in zebrafish *Danio rerio* (and vertebrates generally). They represent major anatomical or developmental domains that emerge during embryogenesis and give rise to specific organs [6]. See also the context of Table S1 that summarizes the major developmental milestones, adapted from the standard Zebrafish staging guide by Kimmel *et al.* [16].

**Central Nervous System.** Composed of the brain and spinal cord. Derived from the neural plate, part of the ectoderm. Includes neurons, glia, and associated structures.

**Endoderm.** One of the three germ layers (along with mesoderm and ectoderm).

Gives rise to gut, liver, pancreas, and lungs (in species that have lungs). Forms internal organs and linings of digestive and respiratory tracts.

**Hematopoietic System.** The system responsible for blood and immune cell formation. Derived mainly from mesodermal tissues. Includes precursors for red blood cells, white blood cells, and other lineages.

**Intermediate Mesoderm.** A part of the mesoderm located between paraxial and lateral mesoderm. Gives rise to kidneys and gonads (urogenital system).

**Lateral Mesoderm.** Forms the circulatory system, body wall, and limb connective tissues. Splits into somatic and splanchnic mesoderm, important for heart and vasculature.

**Mesenchyme.** A loosely organized, migratory cell population derived from multiple germ layers. Gives rise to connective tissue, bone, cartilage, and some organs. Often used as a general label for undifferentiated mesodermal cells.

**Neural Crest.** A multi-potent, migratory cell population arising at the border of neural and non-neural ectoderm. Gives rise to peripheral neurons, glia, pigment cells, craniofacial cartilage, and more. Unique to vertebrates and highly plastic.

**Notochord.** A midline mesodermal structure essential for patterning the neural tube and somites. It signals to the overlying ectoderm to induce neural plate formation. It later regresses, with remnants contributing to the inter-vertebral discs.

**Paraxial Mesoderm.** Gives rise to somites, which further differentiate into: sclerotome (bones), myotome (muscles), and dermatome (dermis). Key for axial skeleton and segmented muscle formation.

**Periderm.** A transient epithelial layer covering the developing embryo. Derived from surface ectoderm. Functions as a protective barrier during early development.

#### Network analysis

We studied GRNs across three distinct developmental stages: early, middle (phylotypic), and late. Each network was characterized based on a range of structural and statistical metrics, Table S2. Below, we explain each of the less intuitive measures and how they vary across stages:

**Clustering Coefficient.** This measures how often a gene's neighbors are also connected to one another. Values decrease slightly from early to middle and then rise in the late stage, suggesting an initial drop in local redundancy followed by stabilization or reemergence of local subnetworks as cells differentiate.

**Modularity.** Reflects how cleanly the network divides into distinct communities, quantified using the Louvain community detection algorithm. The early and middle networks show comparable modularity, while the late network exhibits a clear decrease. This points to a more distributed, less compartmentalized regulatory structure later in development.

**Between-module weight fraction.** Captures how much interaction crosses community boundaries. This fraction is highest in early development, drops markedly in the phylotypic stage, and slightly rises post-phylotypically. The dip at mid-development suggests reduced cross-talk and stronger intra-module cohesion, consistent with maximal developmental constraint.

**Assortativity.** All three networks are negatively assortative, meaning high-degree nodes (hubs) tend to interact with low-degree nodes. The least negative value occurs during the phylotypic stage, indicating a slightly more balanced interaction structure.

**Power-law $\alpha$ and $D$.** The $\alpha$ value reflects how fat-tailed the degree distribution is. Lower $\alpha$ means more hubs. The phylotypic network has the highest $\alpha$ (i.e., least heavy-tailed), hinting at a peak in regulatory centralization. $D$ (Kolmogorov-smirnov distance): Goodness-of-fit to the power-law. The lower the better fit.

These findings align with and refine existing models of how GRN complexity evolves during embryogenesis. In early development, regulatory interactions are largely linear and hierarchical, shaped by maternal inputs and early zygotic activation. GRNs at this stage are sparse and centrally organized, guiding foundational patterning events such as germ layer formation and axis specification. During the phylotypic period, signaling pathways (e.g., Hox, Wnt, FGF, Notch) converge to coordinate the emergence of conserved body structures. Although overall network density increases, this stage is marked by tightly modular architecture with reduced cross-module connectivity, limited local redundancy, and enhanced centralization of regulatory influence; features indicative of global coordination under strong developmental constraint. In later stages, GRNs reorganize toward more distributed, tissue-specific programs. Connectivity becomes more redundant and less modular, reflecting functional diversification and physiological maturation.

#### Perturbation analyses

We implemented five classes of perturbation analyses to assess whether learned embedding representations exhibit heightened sensitivity at the phylotypic stage. These analyses progress from isolated single-gene perturbations to coordinated multi-gene perturbations, include eligibility-matched designs that control for stage-specific gene availability, and extend to topology-aware perturbations that explicitly respect gene interaction structure.

**Single-gene perturbations.** To assess sensitivity to isolated gene removal, we first constructed a high-coverage gene panel shared across developmental stages. From each stage (pre-phylotypic, phylotypic, post-phylotypic), 1,000 cells were randomly sampled. For each sampled cell, gene presence was recorded, and presence counts were pooled across stages. Genes were ranked by the number of sampled cells in which they appeared, and the top 500 genes were selected. This procedure favors genes that are broadly present across cells and stages, yielding a high-coverage but non-stage-specific panel.

For each developmental stage, 300 cells were randomly sampled. Within each cell, up to 10 panel genes present in that cell were selected at random and perturbed *one at a time* by replacing the corresponding token with the ⟨`mask`⟩ token, while preserving sequence length and all other gene positions. For each perturbation, the model was run on the original and masked sequences, hidden-state embeddings were extracted from the final transformer layer, and cosine distances were computed for all positions except the masked one. Distances were averaged across positions to quantify the embedding-space impact of removing a single gene. Per-cell sensitivity scores were obtained by averaging across the 10 perturbations within each cell.

To estimate a numerical baseline, null perturbations were computed on approximately 100 cells per stage by running the model twice on identical unperturbed inputs and computing cosine distances in the same manner. Across three random seeds, single-gene masking induced small but reproducible embedding shifts that exceeded null baselines but were nearly identical across stages, providing no evidence for a phylotypic peak in sensitivity (Table S3).

**Perturbations targeting phylotypic-enriched transcription factors.** Because the high-coverage panel is expected to be dominated by broadly expressed genes, we next focused on transcription factors (TFs) preferentially present at the phylotypic stage (TF list from [17]). For each TF, prevalence was computed separately in pre-phylotypic, phylotypic, and post-phylotypic cells as the fraction of cells in which the TF appeared. A phylotypic enrichment score was defined as the difference between phylotypic prevalence and the maximum of pre- or post-phylotypic prevalence. TFs with phylotypic prevalence below 5% were excluded, yielding a panel of 234 *phylotypic-enriched* TFs.

For each stage, 300 cells were sampled. In each cell, up to 10 TFs from the panel that were present were masked one at a time using the same protocol as above. In addition to the mean embedding shift, two tail-sensitive metrics were computed: the mean of the top 20 largest shifts (top-$k$ mean) and the 99th percentile of shifts ($q_{0.99}$). Per-cell scores were obtained by averaging across masked TFs, and null baselines were estimated as before.

Across three random seeds, targeted masking of phylotypic-enriched TFs produced embedding-space shifts that exceeded numerical baselines but remained similar in magnitude across developmental stages for all metrics (Table S4). Mean and median values were closely matched, indicating tight per-cell distributions and no evidence for rare catastrophic responses.

**Multi-hit perturbations.** To test whether *coordinated* regulatory disruption reveals stage-specific fragility, we next performed multi-hit perturbations in which multiple TFs were masked simultaneously. For each cell, if at least $K = 10$ phylotypic-enriched TFs were present, $K$ were selected at random and all corresponding tokens were replaced with ⟨`mask`⟩ in a single forward pass. Embedding changes were quantified using the same mean, top-$k$, and $q_{0.99}$ metrics, as above, excluding the masked positions. Null baselines were computed by repeated forward passes on identical inputs.

Multi-hit perturbations produced embedding shifts that were two orders of magnitude larger than null baselines, confirming that coordinated perturbations strongly disrupt embedding representations (Table S5). However, effect sizes remained similar across stages, with no consistent elevation at the phylotypic stage. Because the TF panel is phylotypic-enriched, substantially fewer post-phylotypic cells satisfied the requirement of containing at least 10 panel TFs, leading to reduced sample sizes and increased variability in the post-phylotypic estimates.

**Eligibility-matched multi-hit perturbations.** To eliminate potential selection bias arising from unequal TF availability across stages, we implemented an *eligibility-matched* multi-hit protocol. Cells were first filtered to those containing at least $K = 10$ phylotypic-enriched TFs. An equal number of eligible cells ($m = 150$) was then randomly sampled from each stage. Multi-hit perturbations and null controls were applied as above.

Under eligibility-matched conditions, multi-hit perturbations again induced large embedding-space shifts well above numerical baselines, but no systematic increase in sensitivity was observed at the phylotypic stage for any metric (Table S6, Fig. S5). Thus, even when perturbation strength and cell composition are explicitly matched across stages, embedding representations do not exhibit heightened phylotypic fragility.

The perturbation analyses above probe robustness of learned embedding representations to local token removal but do not explicitly account for the interaction structure among genes. To assess whether developmental-stage-specific sensitivity emerges when perturbations respect regulatory topology, we implemented a *topology-aware* perturbation scheme based on attention-derived GRNs.

**Topology-aware perturbations.** For each developmental stage, gene-gene interaction networks were constructed from model attention weights by aggregating pairwise attention scores across cells. Within each stage-specific network, genes were ranked by degree, and high-degree nodes were identified as candidate regulatory 'hubs'. For each hub gene, we defined a local neighborhood consisting of the hub and its directly connected neighbors, with neighborhood size fixed across stages.

Topology-aware perturbations were performed by simultaneously masking all genes within a hub-centered neighborhood, thereby perturbing a local interaction module rather than an isolated gene. To control for trivial effects of gene number and degree, each hub-centered perturbation was compared to degree-matched random gene sets of identical size drawn from the same stage. Embedding-space sensitivity was quantified as the mean cosine-distance shift across non-masked positions, following the same evaluation protocol used for multi-hit perturbations. Across developmental stages, hub-centered neighborhood perturbations induced substantially larger embedding shifts than degree-matched random controls, confirming that the learned representations are sensitive to structured, topology-respecting perturbations (Fig.5, main text).

Across this hierarchy of perturbation analyses, we find a clear separation between robustness in embedding space and sensitivity to interaction structure. Isolated single-gene masking, targeted perturbations of phylotypic-enriched TFs, and increasingly strong multi-hit perturbations all induce reproducible embedding-space shifts that substantially exceed numerical baselines. However, in all cases, these effects remain comparable across developmental stages, even under eligibility-matched multi-hit perturbations that explicitly control for both perturbation strength and cell composition. These results indicate that the learned embedding representations are robust to substantial local and combinatorial gene perturbations throughout development, and that phylotypic-stage specialization is not expressed as increased fragility in embedding space.

In contrast, perturbations that explicitly respect interaction topology reveal sensitivity to network structure that is not captured by token-level embedding robustness. Topology-aware perturbations targeting hub-centered interaction neighborhoods produce substantially larger and more structured embedding responses than degree-matched random perturbations, demonstrating that the model encodes biologically meaningful interaction organization.

In the *pre-phylotypic* stage, overall connectivity is lower and the network is more hierarchical, but hub neighborhoods are relatively self-contained; perturbing them disrupts core early regulators with minimal redundancy. In the *phylotypic* network, connectivity is denser and modularity remains high, but crucially, between-module connectivity is minimal, meaning that perturbing a hub-centered neighborhood still largely affects only a single, insulated module. As a result, both stages show comparable levels of sensitivity to neighborhood ablations, though for slightly different structural reasons: early networks because they are sparse and top-heavy, phylotypic networks because they are dense but modularly compartmentalized. In contrast, *post-phylotypic* GRNs exhibit a marked drop in modularity and a rise in cross-module connectivity, along with more distributed and redundant local structure. Hubs in this regime are less central, and their neighborhoods are embedded in overlapping communities. Perturbing a hub region thus tends to disperse rather than concentrate its effect, and the system absorbs the damage more effectively. This buffering, combined with the flatter degree distribution, explains the reduced perturbation impact observed in this stage, despite its higher raw connectivity (Fig.5, main text).

#### Functional enrichment analysis of attention across development

To characterize the functional roles of genes within attention-derived GRNs, we analyzed directed interaction matrices constructed from normalized and sparsified attention scores. In these matrices, rows correspond to genes exerting influence (outgoing attention), and columns correspond to genes receiving influence (incoming attention). For each developmental stage, we quantified gene influence using weighted out-strength (influencing) and in-strength (influenced). Genes in the top 10th percentile of each distribution were classified as influencing, influenced, or common (high in both). Gene Ontology (GO) enrichment analysis was performed using GProfiler (https://biit.cs.ut.ee/gprofiler/gost) via its Python interface, restricted to molecular function terms and using the expressed gene set as background (Table S7).

`Early Stage.` Genes with high in-strength were strongly enriched for RNA binding, nucleic acid binding, chromatin binding, and nucleosome-associated functions, indicating that early development is characterized by coordinated regulation of transcriptional and post-transcriptional machinery. In contrast, no strong molecular-function enrichment was observed among influencing genes, suggesting heterogeneous upstream regulatory inputs rather than a unified class of molecular drivers.

`Middle (Phylotypic) Stage.` The phylotypic stage showed pronounced enrichment of chromatin- and DNA-associated molecular functions among influenced genes, including chromatin binding, nucleosome binding, and structural constituents of chromatin. This indicates convergence of regulatory attention onto a conserved chromatin-centered core. Influencing genes exhibited weaker but specific enrichment for signaling-related functions, consistent with multiple upstream cues coordinating a shared regulatory architecture.

`Late Stage.` In late development, influencing genes were strongly enriched for signaling receptor activity, transmembrane signaling, and extracellular structural functions, reflecting specialized cell–cell communication and tissue-specific signaling. Influenced genes, in contrast, were enriched for structural, ribosomal, and metabolic molecular functions, consistent with execution of differentiated cellular programs.

Overall, this refined analysis reveals a progressive shift from heterogeneous regulatory inputs in early development, to centralized chromatin-focused regulation at the phylotypic stage, and finally to specialized signaling-driven control of differentiated cellular machinery. These results support an hourglass-like organization of regulatory influence, while clarifying that mid-developmental constraint arises from convergence onto a conserved regulatory core rather than from uniform global interactivity.

#### The transformer architecture in large language models

Transformers are a fundamental multilayer neural architecture at the core of modern large language models (LLMs). Their primary function is to map sequences of `input` vectors – representing basic language units or *tokens*– to sequences of `output` vectors that incorporate contextual information into the initial representations [18]. This effectively puts the original notion of the *distributional hypothesis* into action: words that occur in similar contexts tend to have similar meanings, e.g., [19]. Context is captured through a mechanism called self-attention (SA).

**The core idea of SA.** We said above that LLMs represent tokens in terms of vectors in a multidimensional space generally known as *embedding* space; the dimension of this space defining the size of the model, is $d_m$. We can ask how similar these representations are for two given tokens. Having vectors, this suggests the scalar dot product as a potentially valid score. Thus, if we consider a token $i$ represented by vector $\vec{x}_i$, we can compute its similarity to any other token $j$ represented by vector $\vec{x}_j$ within the LLM *vocabulary* (the total set of unique tokens that the model can recognize and process) as

$$\alpha_{ij} = \frac{\exp(\vec{x}_i \cdot \vec{x}_j)}{\sum_{k=1}^{i} \exp(\vec{x}_i \cdot \vec{x}_k)} = \text{softmax}(\vec{x}_i \cdot \vec{x}_j). \tag{1}$$

This equation represents a *normalized* similarity between tokens $i$ and $j$ with $\alpha_{ij}$ quantifying how much the representation at position $i$ should "pay attention" to (i.e., be influenced by) the token at position $j$. The value is higher when $\vec{x}_i$ and $\vec{x}_j$ are more similar. Note also that Eqn. (1) is limited to preceding words only on *causal* context (backward-looking SA) rather than *bidirectional* context, which incorporates future words. The latter will be discussed in the following sections. The output after the attention is given by $\vec{a}_i = \sum_{j \le i} \alpha_{ij}\vec{x}_j$, a linear combination of vectors $\vec{x}_j$ (Figure S6).

**The attention head.** The attention head is the version of attention used in transformers. The jargon "head" often refers to parallel processing units that operate on the same input but capture different features. A novel notion here is that now we are going to use the input vectors in three different manners, as *query* (the vector that asks, "who is relevant to me?"), as *key* (the vector that answers, "am I relevant to this query?", and as value (the actual content that gets passed to the next layer, weighted by attention scores). This terminology has its roots in information retrieval and search systems, see also [20].

Thus, each word (or token) in the sequence is associated with three special vectors: the *query* $\vec{q}_i$, the *key* $\vec{k}_i$, and the *value* $\vec{v}_i$. These vectors are computed by applying distinct learned weight matrices to the original input vector (embedding) $\vec{x}_i$, specifically using the matrices $W^{\mathtt{Q}}$, $W^{\mathtt{K}}$, and $W^{\mathtt{V}}$. But what does "learned" mean in this context? In machine learning terminology, this means that the values of these matrices are *not fixed in advance.* Instead, they are initialized randomly and then *updated gradually during training.*

As the model is exposed to large amounts of data, it updates these weights to reduce prediction error, effectively learning useful representations that help capture meaningful relationships between tokens. Each weight matrix projects the input vector $\vec{x}_i$ into a different subspace, emphasizing different aspects of its role in context (Figure S7):

$$\begin{aligned} \vec{q}_i &= \vec{x}_i W^{\mathtt{Q}}, \\ \vec{k}_i &= \vec{x}_i W^{\mathtt{K}}, \\ \vec{v}_i &= \vec{x}_i W^{\mathtt{V}}. \end{aligned} \tag{2}$$

Specifically, $W^{\mathtt{Q}}, W^{\mathtt{K}} \in \mathbb{R}^{d_m \times d_k}$ and $W^{\mathtt{V}} \in \mathbb{R}^{d_m \times d_v}$, where $d_m$ is the size of the input embedding, as mentioned before, and $d_k$, $d_v$ are the dimensions of the query/key and value vectors, respectively. These transformed vectors are then used to compute the SA output vector $\vec{a}_i$, which captures how strongly the current token attends to others in the sequence. The new set of equations for computing SA for a single output vector $\vec{a}_i$ from a single input vector $\vec{x}_i$ now reads:

$$\begin{aligned}\alpha_{ij} &= \text{softmax}(\frac{\vec{q}_i \cdot \vec{k}_j}{\sqrt{d_k}}), \\ \vec{a}_i &= \sum_{j \le i} \alpha_{ij} \vec{v}_j, \\ \vec{h}_i &= \text{head}(\vec{x}_i) = \vec{a}_i W^{\circ},\end{aligned} \tag{3}$$

where $W^{\circ}$ here is a matrix of dimension $[d_{\text{v}} \times d_{\text{model}}]$ so that the dimension of the output embedding is that of the input.

**Multihead attention.** Is a single "head" enough? Different contexts may require different perspectives on how tokens relate to one another, so a single attention mechanism might struggle to capture all relationships effectively. Transformers employ a mechanism where *multiple* attention heads operate in parallel at the *same* layer depth within the model. Each head learns a different projection of the input data, allowing the model to capture diverse relationships simultaneously. Each attention head has its own set of parameters, enabling it to focus on different aspects of the input sequence and set of $W^{\mathtt{Q}}, W^{\mathtt{K}}, W^{\mathtt{V}}$ matrices. The multihead reads then as:

$$\text{MultiHeadAttention}(\vec{x}_i) = (\text{head}_1 \oplus \ldots \oplus \text{head}_h)(\vec{x}_i). \tag{4}$$

Note that $W^{\circ}$ here is now dimension $[hd_{\text{v}} \times d_{\text{model}}]$, with $h$ being the number of heads.

**The transformer `block`.** Assuming the intuition behind SA is now clear, what remains is to understand the three additional components of the transformer *block*. We encounter the usual suspects typical of machine learning: issues with learning efficiency and stability.

We then introduce three additional components that help address these issues. (i) A Feedforward Network (FFN): A small, fully connected network applied independently to each position. This network introduces *non-linearity* and helps transform the representations learned from SA. (ii) Residual Connections: These connections allow the original input to be added back to the transformed output at different stages. This technique helps *stabilize* training, prevent vanishing gradients –a known problem that happens when learning signals become too small in deep networks, making it hard for early layers to learn– and ensure efficient gradient propagation (this is associated with the main learning algorithm, backpropagation), and (iii) Normalization Layers (Layer normalize): These layers standardize the inputs to improve numerical *stability* and ensure better generalization. Normalization helps prevent issues like internal covariate shift, where different parts of the network see inputs with drastically different scales, making the network more robust.

Thus, each input vector $\vec{x}_i$ (representing a token at position $i$) is processed through the following steps within a transformer block (*tb*, we skip vector notations for simplicity, Figure S8):

$$
\begin{aligned}
tb_1^i &= \text{LayerNorm}(x_i), \\
tb_2^i &= \text{MultiHeadAttention}(tb_1^i), \\
tb_3^i &= tb_2^i + x_i, \\
tb_4^i &= \text{LayerNorm}(tb_3^i), \\
tb_5^i &= \text{FFN}(tb_4^i), \\
h^i &= tb_5^i + tb_3^i. \qquad (5)
\end{aligned}
$$

In simpler terms, the input is first normalized to stabilize and speed up training by maintaining consistent value scales ($tb_1^i$). SA is applied, allowing the token to incorporate information from other tokens in the sequence ($tb_2^i$). The attention output is added back to the original input to preserve information and aid gradient flow ($tb_3^i$). The combined output is normalized again before further processing ($tb_4^i$). A position-wise FFN further transforms the representation, typically using two linear layers with a non-linear activation ($tb_5^i$). The output of the FFN is added to its input, producing the final output $\vec{h^i}$ of the transformer block for token $i$ (Figure S8).

Finally, note that for simplicity, this introduction describes the transformer as a causal, or left-to-right, language model –the model reads text one word at a time and predicts the next word. However, Zebraformer is a bidirectional transformer encoder, similar to the BERT model [21–23]. This type of model is trained using masked language modeling: instead of guessing the next word, it learns to fill in a missing word somewhere in the middle of a sentence, using clues from both the left and right sides. This approach allows the model to understand a word based on its full context, unlike the left-to-right models shown in Fig. S6.

This section has provided only a brief introduction to the rich and complex world of transformers. A thoughtful overview of language processing can be found in [24], which served as the basis for these notes. There are plenty of great texts and online resources where you can actually learn to program transformers, which is, honestly, the best way to really understand them. You might also wonder how such ideas came about –could *you* have invented transformers? One very interesting perspective on that question can be found here: https://gwern.net/blog/2025/you-could-have-invented-transformers.

#### LLMs and single-cell data

In some areas of biology, applying LLMs is a natural fit because the concepts of *sequence* and *context* are central to the data. As discussed in the previous section, LLMs learn patterns based on the relationships between elements in a sequence. The information is derived not from isolated components, but from their surrounding context. This makes LLMs especially well-suited to problems like protein modeling, where the linear arrangement of amino acids determines structure and function. Indeed, LLMs have been successfully applied to protein sequences to predict structure, function, and evolutionary relationships, e.g., [25–27]. Similar strategies are also gaining traction in genomics and RNA biology, where biological meaning often depends on the position and context of sequence elements.

In contrast, single-cell transcriptomic data presents an initial challenge: it is not inherently sequential. Rather than representing a chain of elements, it captures the activity (expression levels) of many genes simultaneously across the genome. To make this data compatible with transformers, which expect sequence-like input, it must first be adapted into a format that imposes a meaningful order. How can we do this? So far, three main approaches have been used: i/ *Ordering*: This method transforms the expression profile of a cell into a sequence by ranking genes according to their expression levels. Each gene becomes a token in the sequence, similar to a word in a sentence. This ranked sequence is then passed to the transformer model. This is the approach adopted in our work (see below); ii/ *Value Categorization*: Gene expression values are grouped into discrete bins, e.g., low, medium, high, and each bin receives a distinct embedding. This makes the continuous expression data compatible with models designed for categorical inputs. iii/ *Value Projection*: Rather than simplifying the data, as in i/ and ii/, this method retains the full continuous expression values and projects them into a new space using mathematical transformations (often linear).

In our work, we build on the architecture and methods introduced in Geneformer [23], a transformer model based on the `BERT` architecture [21], see also [22]. Recall from the previous section that unlike *causal* language models, `BERT` uses bidirectional attention, allowing it to consider the full context surrounding each token simultaneously; an advantage when dealing with non-linguistic data like gene expression. Geneformer employs a rank-based encoding scheme in which genes are ordered by their normalized expression levels within each cell. By trying to predict missing (masked) genes in a sequence based on the surrounding context the model learns contextual relationships between genes and builds rich internal representations (embeddings) of them (see Methods, main text).

Finally, to help situate our work in the broader landscape of LLMs for single-cell omics, we include a summary table (Table S8) highlighting a selection of some recent models. This is not intended to be an exhaustive list, but rather a snapshot of current trends (see also references on the main text; we apologize for any important omissions). For a more comprehensive review, see [28].

### References


1. Karl Ernst von Baer. *Über Entwickelungsgeschichte der Thiere: Beobachtung und Reflektion*. Koenigsberg, Koenigsberg, 1828.

2. Ernst Haeckel. *Generelle Morphologie der Organismen. Allgemeine Grundzüge der organischen Formen-Wissenschaft, mechanisch begründet durch die von Charles Darwin reformirte Descendenz-Theorie*. Georg Reimer, Berlin, 1866.

3. Jonathan MW Slack, Peter WH Holland, and Charles F Graham. The zootype and the phylotypic stage. *Nature*, 361:490–492, 1993.

4. Denis Duboule. Temporal colinearity and the phylotypic progression: a basis for the stability of a vertebrate Bauplan and the evolution of morphologies through heterochrony. *Development: Supplement*, pages 135–142, 1994.

5. Rudolf A. Raff. *The Shape of Life: Genes, Development, and the Evolution of Animal Form.* University of Chicago Press, Chicago, IL, 2000.

6. Lewis Wolpert, Cheryll Tickle, Alfonso Martinez Arias, Peter Lawrence, and James Locke. *Principles of Development.* Oxford University Press, Oxford, UK, 6th edition, 2023.

7. Eric H. Davidson. *The Regulatory Genome: Gene Regulatory Networks in Development and Evolution.* Academic Press, Burlington, MA, 2nd edition, 2006.

8. Frietson Galis and Johan A. J. Metz. Testing the vulnerability of the phylotypic stage: on modularity and evolutionary conservation. *Proceedings: Biological Sciences*, 268(1477):181–188, 2001.

9. Naoki Irie and Shigeru Kuratani. The developmental hourglass model: a predictor of the basic body plan? *Development*, 141(24):4649–4655, 2014.

10. Merlin Lange, Alejandro Granados, Shruthi VijayKumar, Jordão Bragantini, Sarah Ancheta, Yang-Joon Kim, Sreejith Santhosh, Michael Borja, Hirofumi Kobayashi, Erin McGeever, Ahmet Can Solak, Bin Yang, Xiang Zhao, Yang Liu, Angela M. Detweiler, Sheryl Paul, Ilan Theodoro, Honey Mekonen, Chris Charlton, Tiger Lao, Rachel Banks, Sheng Xiao, Adrian Jacobo, Keir Balla, Kyle Awayan, Samuel D'Souza, Robert Haase, Alexandre Dizeux, Olivier Pourquie, Rafael Gómez-Sjöberg, Greg Huber, Mattia Serra, Norma Neff, Angela Oliveira Pisco, and Loïc A. Royer. A multimodal zebrafish developmental atlas reveals the state-transition dynamics of late-vertebrate pluripotent axial progenitors. *Cell*, 187(23):6742–6759.e17, 2024.

11. Daniel E. Wagner, Caleb Weinreb, Zach M. Collins, James A. Briggs, Sean G. Megason, and Allon M. Klein. Single-cell mapping of gene expression landscapes and lineage in the zebrafish embryo. *Science*, 360(6392):981–987, 2018.

12. Jeffrey A. Farrell, Yiqun Wang, Samantha J. Riesenfeld, Karthik Shekhar, Aviv Regev, and Alexander F. Schier. Single-cell reconstruction of developmental trajectories during zebrafish embryogenesis. *Science*, 360(6392):eaar3131, 2018.

13. Dylan R. Farnsworth, Lauren M. Saunders, and Adam C. Miller. A single-cell transcriptome atlas for zebrafish development. *Developmental Biology*, 459(2):100–108, 2020.

14. Lauren M. Saunders, Sanjay R. Srivatsan, Madeleine Duran, Michael W. Dorrity, Brent Ewing, Tor Linbo, Jay Shendure, David W. Raible, Cecilia B. Moens, David

Kimelman, and Cole Trapnell. Deep molecular, cellular and temporal phenotyping of developmental perturbations at whole organism scale, 2022.

15. Abhinav Sur, Yiqun Wang, Paulina Capar, Gennady Margolin, Morgan Kathleen Prochaska, and Jeffrey A. Farrell. Single-cell analysis of shared signatures and transcriptional diversity during zebrafish development. *Developmental Cell*, 58(24):3028–3047.e12, 2023.

16. Charles B. Kimmel, William W. Ballard, Sarah R. Kimmel, Bonnie Ullmann, and Thomas F. Schilling. Stages of embryonic development of the zebrafish. *Developmental Dynamics*, 203(3):253–310, 1995.

17. Wen-Kang Shen, Si-Yi Chen, Zi-Quan Gan, Yu-Zhu Zhang, Tao Yue, Miao-Miao Chen, Xue Yu, Hui Hu, and An-Yuan Guo. AnimalTFDB 4.0: a comprehensive animal transcription factor database updated with variation and expression annotations. *Nucleic Acids Research*, 51(D1):D39–D45, 2023.

18. Ashish Vaswani, Noam Shazeer, Niki Parmar, Jakob Uszkoreit, Llion Jones, Aidan N. Gomez, Łukasz Kaiser, and Illia Polosukhin. Attention is all you need. In *Advances in Neural Information Processing Systems (NeurIPS)*, volume 30, pages 5998–6008. Curran Associates, Inc., 2017.

19. Martin Joos. Description of language design. *The Journal of the Acoustical Society of America*, 22(6):701–708, 1950.

20. Dzmitry Bahdanau, Kyunghyun Cho, and Yoshua Bengio. Neural machine translation by jointly learning to align and translate. In *Proceedings of the 3rd International Conference on Learning Representations (ICLR)*, 2014.

21. Jacob Devlin, Ming-Wei Chang, Kenton Lee, and Kristina Toutanova. BERT: Pre-training of deep bidirectional transformers for language understanding. In *Proceedings of the 2019 Conference of the North American Chapter of the Association for Computational Linguistics: Human Language Technologies, Vol. 1*, pages 4171–4186. Association for Computational Linguistics, 2019.

22. Fan Yang, Wenchuan Wang, Fang Wang, Yuan Fang, Duyu Tang, Junzhou Huang, Hui Lu, and Jianhua Yao. scBERT as a large-scale pretrained deep language model for cell type annotation of single-cell rna-seq data. *Nature Machine Intelligence*, 4(10):852–866, 2022.

23. Christina V. Theodoris, Ling Xiao, Anant Chopra, Mark D. Chaffin, Zeina R. Al Sayed, Matthew C. Hill, Helene Mantineo, Elizabeth M. Brydon, Zexian Zeng, X. Shirley Liu, and Patrick T. Ellinor. Transfer learning enables predictions in network biology. *Nature*, 618(7965):616–624, 2023.

24. Daniel Jurafsky and James H. Martin. *Speech and Language Processing: An Introduction to Natural Language Processing, Computational Linguistics, and Speech Recognition with Language Models.* 3rd edition, 2025.

25. Alexander Rives, Joshua Meier, Tom Sercu, Siddharth Goyal, Zeming Lin, Jason Liu, Demi Guo, Myle Ott, C Lawrence Zitnick, Jerry Ma, et al. Biological structure and function emerge from scaling unsupervised learning to 250 million protein sequences. *Proceedings of the National Academy of Sciences*, 118(15), 2021.

26. Nadav Brandes, Dan Ofer, Yam Peleg, Nadav Rappoport, and Michal Linial. Proteinbert: A universal deep-learning model of protein sequence and function. *Bioinformatics*, 38(8):2102–2110, 2022.

27. Zeming Lin, Halil Akin, Roshan Rao, Brian Hie, Zhongkai Zhu, Wenting Lu, Nikita Smetanin, Robert Verkuil, Ori Kabeli, Yaniv Shmueli, et al. Evolutionary-scale prediction of atomic-level protein structure with a language model. *Science*, 379(6637):1123–1130, 2023.

28. Artur Szałata, Karin Hrovatin, Sören Becker, Alejandro Tejada-Lapuerta, Haotian Cui, Bo Wang, and Fabian J. Theis. Transformers in single-cell omics: a review and new perspectives. *Nature Methods*, 21(8):1430–1443, 2024.

29. Jing Gong, Minsheng Hao, Xingyi Cheng, Xin Zeng, Chiming Liu, Jianzhu Ma, Xuegong Zhang, Taifeng Wang, and Le Song. xTrimoGene: An efficient and scalable representation learner for single-cell RNA-seq data. *arXiv preprint arXiv:2311.15156*, 2023.

30. Suyuan Zhao, Jiahuan Zhang, Yushuai Wu, Yizhen Luo, and Zaiqing Nie. LangCell: Language-cell pre-training for cell identity understanding. *arXiv preprint arXiv:2405.06708*, 2024.

31. Moritz Schaefer, Peter Peneder, Daniel Malzl, Mihaela Peycheva, Jake Burton, Anna Hakobyan, Varun Sharma, Thomas Krausgruber, Jörg Menche, Eleni M. Tomazou, and Christoph Bock. Multimodal learning of transcriptomes and text enables interactive single-cell rna-seq data exploration with natural-language chats. *bioRxiv 10.1101/2024.10.15.618501*, 2024.

32. Keita Ito, Tsubasa Hirakawa, Shuji Shigenobu, Hironobu Fujiyoshi, and Takayoshi Yamashita. Mouse-geneformer: A deep learning model for mouse single-cell transcriptome and its cross-species utility. *PLoS Genetics*, 21(3):e1011420, 2025.

33. Cong Qi, Hanzhang Fang, Tianxing Hu, Siqi Jiang, and Wei Zhi. GeneMamba: Efficient and effective foundation model on single cell data. *arXiv preprint arXiv:2504.16956*, 2025.

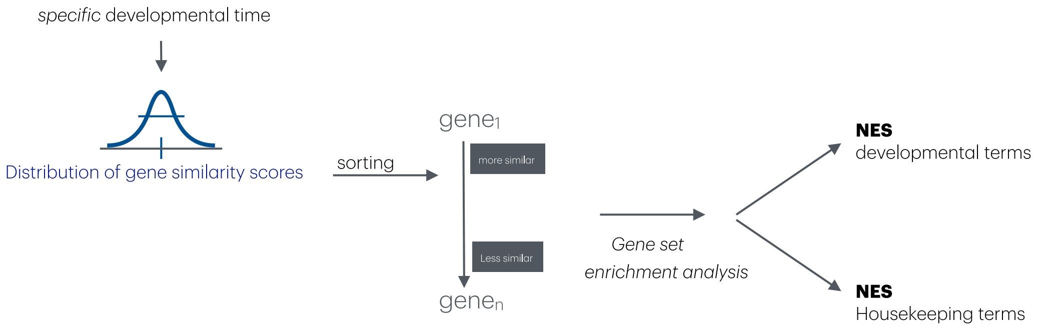


**Figure S1: Normalized enrichment scores.** Given a distribution of gene similarity scores at a specific developmental timepoint, we developed a functional enrichment protocol that first sorts the similarity scores and then computes a normalized enrichment score (NES) associated with either developmental terms ("development", "morphogenesis", "differentiation", "patterning", and "organogenesis") or housekeeping terms ("transcription", "translation", "DNA replication", "metabolism", and "RNA processing").

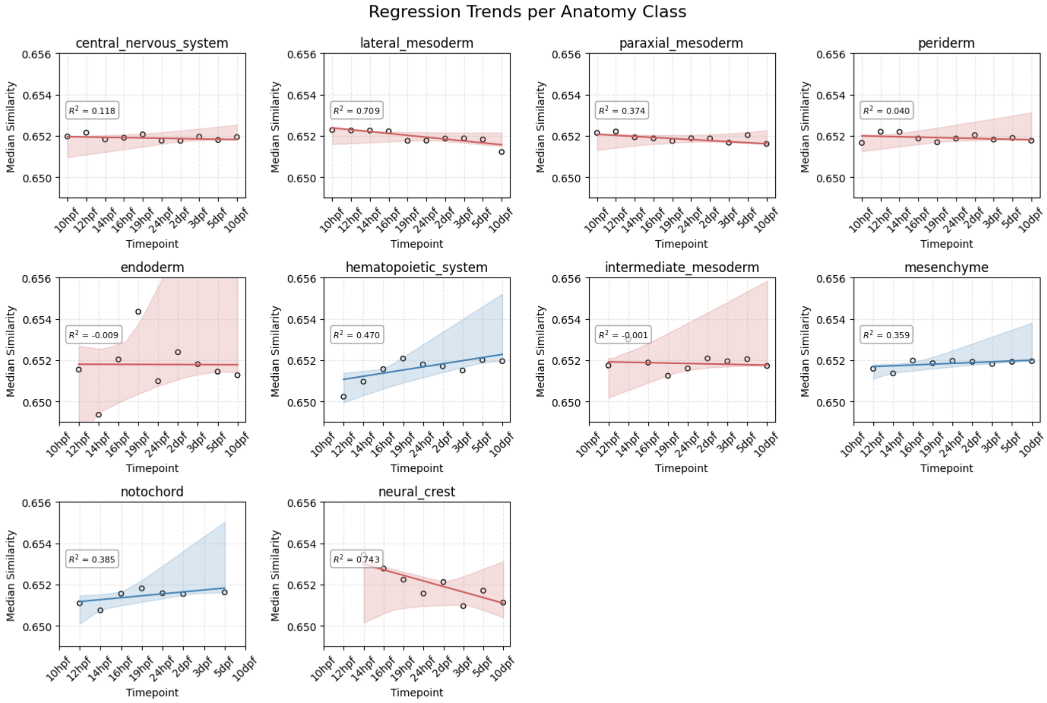


**Figure S2: Gene embeddings and context across all anatomical domains.** Regression trends of median gene embedding similarity for each anatomical class, with shaded regions indicating bootstrapped uncertainty (10,000 samples). For each gene, similarity scores are computed across cells in which the gene is active; the median similarity of all such genes within a given anatomical domain and timepoint reflects the overall contextual consistency of gene representations. Main text for details.

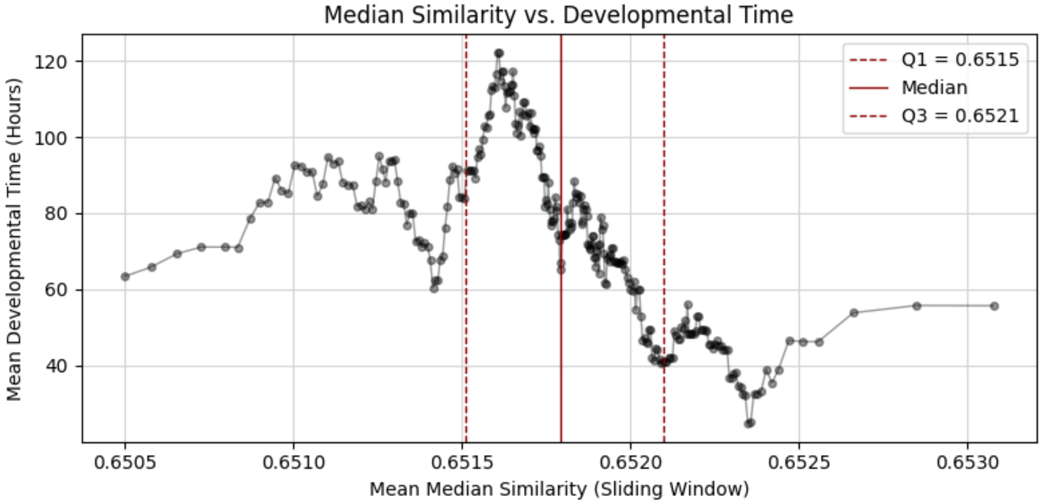


**Figure S3: Gradient of contextual diversity.** Sliding window analysis (window size of 30 cells) of the median similarity scores for genes active on each cell type. The $x$-axis shows the average similarity within each window, and the $y$-axis shows the average corresponding timepoints. The plot also includes the median and the 25th ($Q1$) and 75th ($Q3$) percentiles of the distribution of median similarities. Cell types whose genes exhibit higher average similarity tend to be associated with earlier developmental stages. Line included to aid visualization.

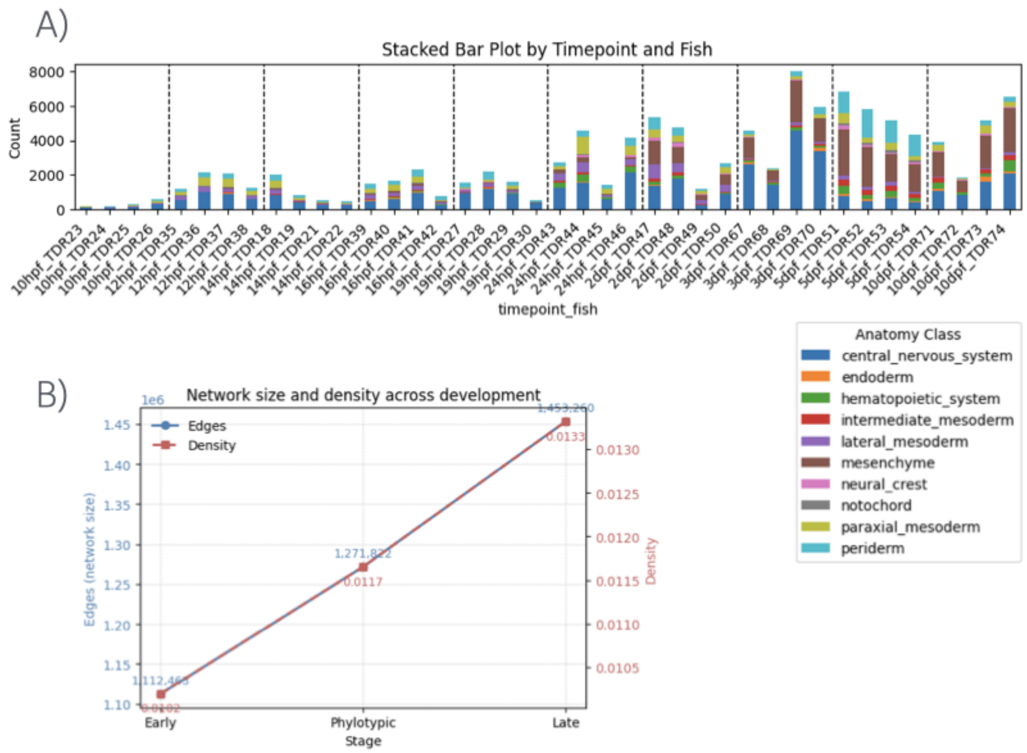


**Figure S4: Anatomical classes by timepoint and fish and change in network main features.** (A) Stacked bar plot showing the distribution of anatomical classes (e.g., zebrafish tissues or structures labeled using zebrahub information; see text) across different fish (TDRxx) and developmental timepoints (10hpf, 12hpf, . . .), reflecting increasing organismal complexity over time. (B) Network size (number of edges, red) and density (blue) increase monotonically from early to late development.

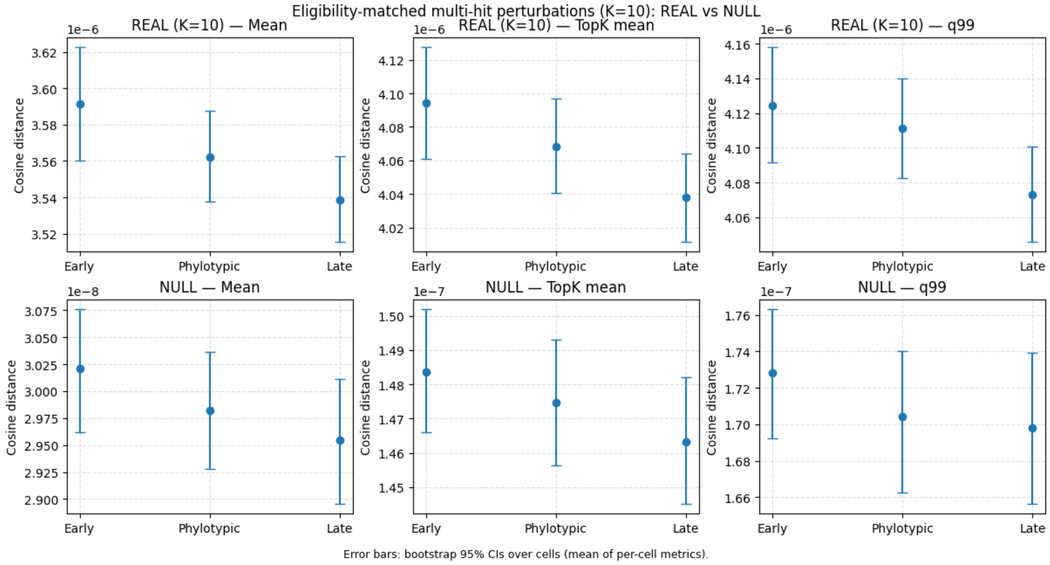


**Figure S5: Eligibility-matched multi-hit perturbations**. Embedding displacement induced by eligibility-matched multi-gene masking ($K = 10$) across early ($\leq$14 hpf), phylotypic (15–19 hpf), and late ($\geq$24 hpf) stages. Top row shows real perturbations; bottom row shows null (random) perturbations. For each stage we report three summaries of per-cell cosine distances between wild-type and perturbed embeddings (Table S6). Points indicate stage averages and error bars denote bootstrap 95% confidence intervals over cells. While embedding displacement decreases modestly from early to late development, this pattern is mirrored by null perturbations indicating a generic property of the embedding geometry.

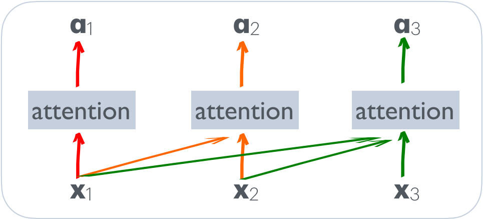


**Figure S6: Causal self-attention (SA).** A transformer processes tokens in a sentence $(\vec{x}_1, \vec{x}_2, \vec{x}_3, \ldots)$ using a SA mechanism. Each token $\vec{x}_i$ attends only to earlier tokens $(j \leq i)$, a setup known as *causal* or *left-to-right* SA. This produces an output vector $\vec{a}_i$ based solely on past context. In contrast, *masked language modeling* (as used in Zebraformer) allows attention to all tokens, both before and after the masked word (vectors shown in bold).

.

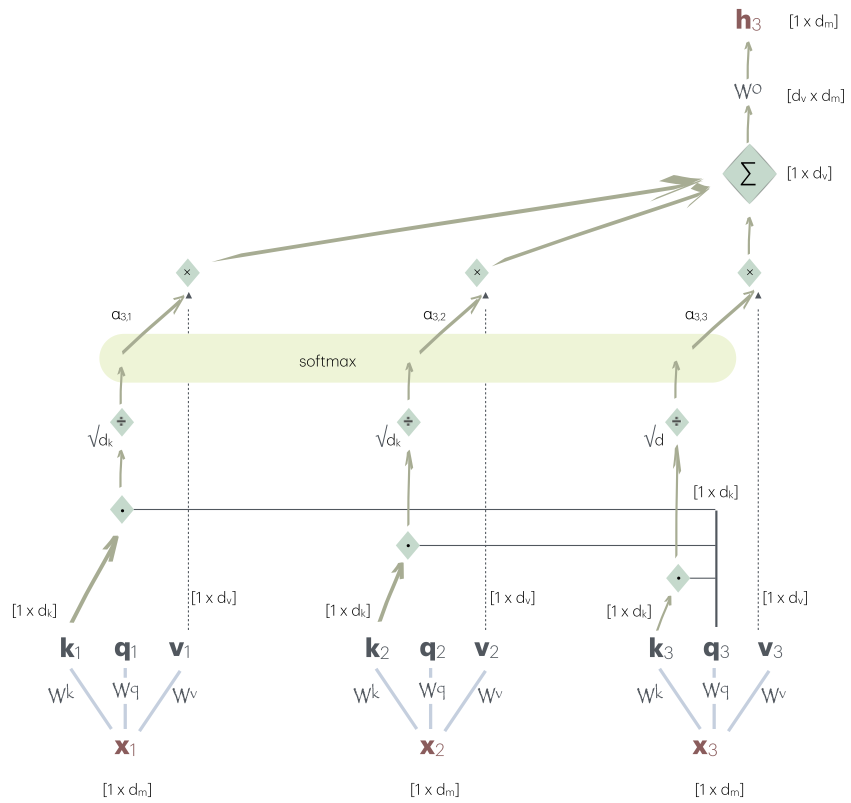


**Figure S7: Self-attention head.** Example of the workings of the attention head for input $\vec{x}_3$ following Eqn. (3), vectors shown in bold. Shown are the three manners –query, key, and value– in which we represent the input vectors and their associated weight matrices. The dot product between queries and keys is scaled by $\sqrt{d_k}$ to prevent large values from pushing the softmax into regions with vanishing gradients, thus stabilizing training. Note the corresponding dimensions at the end and the need for matrix $\mathtt{W}^o$ of dimensions $[d_\mathrm{v} \times d_\mathrm{m}]$. The resultant output vector is $\vec{h}_3$.

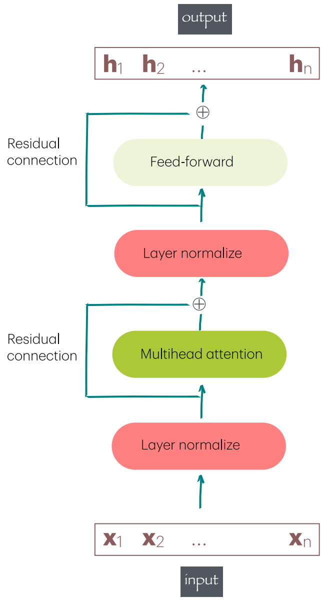


**Figure S8: Computation flow within a transformer block**. Each element of the input sequence $(\vec{x}_1, \vec{x}_2, \vec{x}_3, \ldots)$ is first normalized, then passed through a self attention layer to gather contextual information from other tokens. The result is added back to the original input (residual connection), normalized again, and processed by a feedforward network. A second residual connection produces the final output for each token in the $(\vec{h}_1, \vec{h}_2, \vec{h}_3, \ldots)$ final sequence.

| Time (hpf) | Stage | Key events |
|---|---|---|
| 0–0.75 | Zygote | Fertilization; single-cell with visible cytoplasm and yolk |
| 0.75–2.25 | Cleavage | Rapid, synchronous cell divisions; no transcription yet (maternal control) |
| 2.25–5.25 | Blastula | 128–1k cells; dome forms; zygotic genome activation (ZGA) begins ($\sim$3 hpf) |
| 5.25–10 | Gastrula | Germ layers form (ectoderm, mesoderm, endoderm); epiboly, involution processes occur |
| 10–24 | Segmentation | Somites form; notochord, neural tube develop; neurulation begins |
| 24–48 | Pharyngula | Body plan established; visible eye and brain regions; tail straightens |
| 48–72 | Early Larva | Circulation functional; pectoral fins form; pigmentation appears; hatching |
| 72–5 dpf | Late Larva | Free-swimming; jaw forms; early behavior (touch response) observed |
| 5–7 dpf | Larva to Juvenile | Major organ systems functional; swim bladder inflates; onset of feeding |
| 1–3 weeks | Juvenile | Rapid growth; pigmentation and fin rays develop |
| 3+ weeks | Adult | Sexual maturation; fully developed body morphology |

**Table S1:** Developmental stages and key events in zebrafish embryogenesis. See [16] for details. hpf: hours post-fertilization. dpf: days post-fertilization.

| Property | Early | Middle | Late |
|---|---|---|---|
| Nodes | 14,775 | 14,775 | 14,775 |
| Edges | 1,112,463 | 1,271,822 | 1,453,260 |
| Density | 0.0102 | 0.0117 | 0.0133 |
| Average Degree | 150.59 | 172.16 | 196.72 |
| Clustering Coefficient | 0.1986 | 0.1614 | 0.1675 |
| Average Path Length | 2.19 | 2.09 | 2.00 |
| Diameter | 4 | 4 | 4 |
| Modularity (mean) | 0.2083 | 0.2077 | 0.1833 |
| Modularity (sd) | 0.0030 | 0.0011 | 0.0028 |
| Between-module weight fraction (mean) | 0.6940 | 0.5619 | 0.5887 |
| Between-module weight fraction (sd) | 0.0248 | 0.0255 | 0.0147 |
| Assortativity | -0.2073 | -0.1845 | -0.2232 |
| Power-Law $\alpha$ | 2.30 | 2.37 | 2.26 |
| Power-Law $D$ | 0.0258 | 0.0221 | 0.0276 |

**Table S2:** Network metrics across the three developmental stages. All networks are based on the same set of 14,775 nodes but differ in interaction density and topology.

| Seed | Stage | Real Mean | Real Median | Null Mean |
|---|---|---|---|---|
| | Pre | $9.2505 \times 10^{-8}$ | $9.2547 \times 10^{-8}$ | $1.4868 \times 10^{-8}$ |
| 1 | Phy | $9.2520 \times 10^{-8}$ | $9.2345 \times 10^{-8}$ | $1.4981 \times 10^{-8}$ |
| | Post | $9.2560 \times 10^{-8}$ | $9.2602 \times 10^{-8}$ | $1.4524 \times 10^{-8}$ |
| | Pre | $9.2286 \times 10^{-8}$ | $9.2228 \times 10^{-8}$ | $1.4871 \times 10^{-8}$ |
| 2 | Phy | $9.2882 \times 10^{-8}$ | $9.2998 \times 10^{-8}$ | $1.4729 \times 10^{-8}$ |
| | Post | $9.2457 \times 10^{-8}$ | $9.2281 \times 10^{-8}$ | $1.4815 \times 10^{-8}$ |
| | Pre | $9.2767 \times 10^{-8}$ | $9.2578 \times 10^{-8}$ | $1.5046 \times 10^{-8}$ |
| 3 | Phy | $9.2788 \times 10^{-8}$ | $9.2944 \times 10^{-8}$ | $1.4614 \times 10^{-8}$ |
| | Post | $9.2552 \times 10^{-8}$ | $9.2648 \times 10^{-8}$ | $1.4112 \times 10^{-8}$ |

**Table S3:** Embedding-space sensitivity under single-gene masking across developmental stages. For each random seed, 300 cells per stage were analyzed, with 10 panel genes masked per cell. Reported are the mean and median per-cell embedding impact for real perturbations, and the mean null impact estimated from repeated forward passes on identical inputs ($n = 100$ null cells per stage).

| Stage | Metric | Real Mean | Null Mean |
|---|---|---|---|
| Pre | Mean | $8.75 \times 10^{-8}$ | $3.00 \times 10^{-8}$ |
| | Top-k Mean | $2.06 \times 10^{-7}$ | $1.48 \times 10^{-7}$ |
| | $q_{0.99}$ | $2.24 \times 10^{-7}$ | $1.70 \times 10^{-7}$ |
| Phy | Mean | $8.74 \times 10^{-8}$ | $3.00 \times 10^{-8}$ |
| | Top-k Mean | $2.05 \times 10^{-7}$ | $1.46 \times 10^{-7}$ |
| | $q_{0.99}$ | $2.23 \times 10^{-7}$ | $1.69 \times 10^{-7}$ |
| Post | Mean | $8.73 \times 10^{-8}$ | $2.98 \times 10^{-8}$ |
| | Top-k Mean | $2.05 \times 10^{-7}$ | $1.47 \times 10^{-7}$ |
| | $q_{0.99}$ | $2.23 \times 10^{-7}$ | $1.71 \times 10^{-7}$ |

**Table S4:** Embedding-space sensitivity under targeted masking of phylotypic-enriched transcription factors. Values report averages across three random seeds. For each seed, 300 cells per stage were analyzed, with up to 10 TFs masked per cell. Reported are mean per-cell perturbation metrics ($k = 20$) and corresponding null baselines.

| Stage | Metric | Real (mean ± SD) | Null (mean ± SD) | $n$ (real/null) |
|---|---|---|---|---|
| Pre | Mean | $3.5717 \times 10^{-6} \pm 7.16 \times 10^{-9}$ | $2.9850 \times 10^{-8} \pm 2.77 \times 10^{-10}$ | 886 / 294 |
| | Top-$k$ mean | $4.0727 \times 10^{-6} \pm 7.24 \times 10^{-9}$ | $1.4735 \times 10^{-7} \pm 1.64 \times 10^{-9}$ | 886 / 294 |
| | $q_{0.99}$ | $4.1081 \times 10^{-6} \pm 5.60 \times 10^{-9}$ | $1.7044 \times 10^{-7} \pm 2.74 \times 10^{-9}$ | 886 / 294 |
| Phy | Mean | $3.5640 \times 10^{-6} \pm 2.96 \times 10^{-9}$ | $3.0372 \times 10^{-8} \pm 2.35 \times 10^{-10}$ | 855 / 287 |
| | Top-$k$ mean | $4.0633 \times 10^{-6} \pm 6.52 \times 10^{-9}$ | $1.4684 \times 10^{-7} \pm 7.45 \times 10^{-10}$ | 855 / 287 |
| | $q_{0.99}$ | $4.0971 \times 10^{-6} \pm 7.07 \times 10^{-9}$ | $1.7170 \times 10^{-7} \pm 1.37 \times 10^{-9}$ | 855 / 287 |
| Post | Mean | $3.5594 \times 10^{-6} \pm 2.41 \times 10^{-8}$ | $2.9555 \times 10^{-8} \pm 2.90 \times 10^{-10}$ | 440 / 146 |
| | Top-$k$ mean | $4.0556 \times 10^{-6} \pm 2.97 \times 10^{-8}$ | $1.4667 \times 10^{-7} \pm 2.38 \times 10^{-9}$ | 440 / 146 |
| | $q_{0.99}$ | $4.0888 \times 10^{-6} \pm 2.94 \times 10^{-8}$ | $1.7291 \times 10^{-7} \pm 3.59 \times 10^{-9}$ | 440 / 146 |

**Table S5:** Embedding-space sensitivity under multi-hit masking of phylotypic-enriched TFs ($K = 10$ masked TFs per cell). For each seed, up to 300 cells per stage were sampled; only cells containing at least $K$ panel TFs were retained. Reported values are the mean across seeds ± SD (seed-to-seed), computed over per-cell metrics ($k = 20$). Null values are computed from repeated forward passes on identical inputs, excluding the same positions.

| Stage | Metric | Real Mean | Null Mean | $n$ (Real Null) |
|---|---|---|---|---|
| Pre | Mean | $3.592 \times 10^{-6}$ | $3.021 \times 10^{-8}$ | 150/100 |
| | Top-$k$ mean | $4.094 \times 10^{-6}$ | $1.484 \times 10^{-7}$ | 150/100 |
| | $q_{0.99}$ | $4.124 \times 10^{-6}$ | $1.728 \times 10^{-7}$ | 150/100 |
| Phylotypic | Mean | $3.562 \times 10^{-6}$ | $2.982 \times 10^{-8}$ | 150/100 |
| | Top-$k$ mean | $4.068 \times 10^{-6}$ | $1.475 \times 10^{-7}$ | 150/100 |
| | $q_{0.99}$ | $4.111 \times 10^{-6}$ | $1.704 \times 10^{-7}$ | 150/100 |
| Post | Mean | $3.539 \times 10^{-6}$ | $2.954 \times 10^{-8}$ | 150/100 |
| | Top-$k$ mean | $4.038 \times 10^{-6}$ | $1.463 \times 10^{-7}$ | 150/100 |
| | $q_{0.99}$ | $4.073 \times 10^{-6}$ | $1.698 \times 10^{-7}$ | 150/100 |

**Table S6:** Eligibility-matched multi-hit perturbations using phylotypic-enriched transcription factors. Cells were restricted to those containing at least $K = 10$ panel TFs, and an equal number of eligible cells ($n = 150$) were sampled per developmental stage. Reported values are per-cell embedding-space sensitivity metrics (mean, top-$k$ mean with $k = 20$, and $q_{0.99}$). Null values are computed from repeated forward passes on identical inputs, excluding the same positions.

| Phase | Group | GO Molecular Function Term | p-value | Size |
|---|---|---|---|---|
| Early | Only Influenced | RNA binding | $1.72 \times 10^{-32}$ | 161 |
| | | nucleic acid binding | $2.05 \times 10^{-24}$ | 335 |
| | | binding | $1.59 \times 10^{-19}$ | 692 |
| | | molecular_function | $6.02 \times 10^{-18}$ | 909 |
| | | mRNA binding | $4.20 \times 10^{-11}$ | 51 |
| | | chromatin binding | $1.43 \times 10^{-9}$ | 56 |
| | | histone binding | $1.69 \times 10^{-5}$ | 27 |
| | | nucleosome binding | $9.43 \times 10^{-5}$ | 22 |
| | | protein binding | $1.54 \times 10^{-4}$ | 239 |
| | | mRNA 3'-UTR binding | $1.75 \times 10^{-4}$ | 19 |
| Middle | Only Influenced | molecular_function | $6.68 \times 10^{-20}$ | 918 |
| | | RNA binding | $3.44 \times 10^{-16}$ | 129 |
| | | binding | $2.00 \times 10^{-15}$ | 674 |
| | | nucleic acid binding | $5.81 \times 10^{-15}$ | 304 |
| | | chromatin binding | $8.33 \times 10^{-15}$ | 65 |
| | | structural molecule activity | $7.10 \times 10^{-12}$ | 93 |
| | | nucleosome binding | $4.04 \times 10^{-10}$ | 29 |
| | | structural constituent of chromatin | $6.79 \times 10^{-10}$ | 44 |
| | | nucleosomal DNA binding | $3.63 \times 10^{-9}$ | 25 |
| | | chromatin DNA binding | $5.10 \times 10^{-8}$ | 28 |
| | Only Influencing | G protein-coupled receptor activity | $3.26 \times 10^{-2}$ | 21 |
| Late | Only Influenced | molecular_function | $2.10 \times 10^{-17}$ | 906 |
| | | structural molecule activity | $6.54 \times 10^{-12}$ | 93 |
| | | binding | $5.39 \times 10^{-10}$ | 645 |
| | | structural constituent of ribosome | $8.10 \times 10^{-6}$ | 37 |
| | | protein binding | $6.36 \times 10^{-5}$ | 241 |
| | | proton transmembrane transporter activity | $1.72 \times 10^{-4}$ | 19 |
| | | unfolded protein binding | $7.05 \times 10^{-4}$ | 15 |
| | | nucleosomal DNA binding | $1.18 \times 10^{-3}$ | 18 |
| | | protein-containing complex binding | $1.21 \times 10^{-3}$ | 56 |
| | | oxidoreductase activity | $1.73 \times 10^{-3}$ | 58 |
| | Only Influencing | acrosin binding | $4.42 \times 10^{-8}$ | 10 |
| | | G protein-coupled receptor activity | $2.79 \times 10^{-6}$ | 28 |
| | | structural constituent of egg coat | $4.47 \times 10^{-6}$ | 8 |
| | | olfactory receptor activity | $2.03 \times 10^{-5}$ | 9 |
| | | odorant binding | $1.67 \times 10^{-4}$ | 8 |
| | | U4 snRNA binding | $3.28 \times 10^{-4}$ | 7 |
| | | signaling receptor activity | $1.78 \times 10^{-3}$ | 60 |
| | | molecular transducer activity | $1.78 \times 10^{-3}$ | 60 |
| | | extracellular matrix structural constituent | $5.12 \times 10^{-3}$ | 9 |
| | | transmembrane signaling receptor activity | $3.15 \times 10^{-2}$ | 33 |

**Table S7:** GO molecular function enrichment of attention-derived influencing and influenced genes across embryonic phases. Gene sets were defined using top 10% in- and out-strength in directed, normalized, sparsified attention networks. Size denotes number of genes associated with each term.

| Model | Year | Description |
|---|---|---|
| xTrimoGene | 2023 | Asymmetric transformer model optimized for large-scale scRNA-seq [29]. |
| LangCell | 2024 | Multimodal model using transcriptomic and text data for zero-shot cell-type recognition [30]. |
| CellWhisperer | 2024 | multimodal machine learning model and software that connects transcriptomes and text for interactive single-cell RNA-seq data analysis [31]. |
| Mouse-Geneformer | 2025 | Mouse-specific Geneformer trained on~21M cells; validated cross-species perturbation [32]. |
| GeneMamba | 2025 | Linear-time, pathway-aware transformer trained on~30M cells [33]. |

**Table S8:** Recent large language models applied to single-cell transcriptomics (2023–2025).
